# Thiol depletion and disruption of proteostasis contribute to the phytotoxicity of juglone

**DOI:** 10.64898/2026.04.11.717897

**Authors:** George W. Meyer, Mearaj A. Shaikh, Frederick Mildenhall, MaKenzie Drowns, Chase T. Hearn, Xiaojin Wang, Chao-Jan Liao, Venkatesh P. Thirumalaikumar, Kranthi Varala, Joshua R. Widhalm

## Abstract

- Juglone is the phytotoxic 1,4-naphthoquinone responsible for the allelopathic effects of black walnut (*Juglans nigra*), yet how plants perceive and respond to juglone remain poorly understood.
- We conducted transcriptome profiling of rosettes and roots of *Arabidopsis thaliana* exposed to juglone from 30 min to 5 d, along with targeted metabolic profiling, biochemical assays, and untargeted proteomics to gain a systems-level understanding of how plants respond to juglone and to test hypotheses underlying its phytotoxicity.
- Juglone exposure induced expression of genes involved in glutathione, cysteine, and sulfur metabolism pathways, and in protein homeostasis. We found that juglone depletes the pool of reduced glutathione (GSH) in roots, in part, through conjugation. We demonstrate that via upregulation of transcription factors (*NAC53* and *NAC78*), the response to juglone activates components of the proteasome stress regulon and triggers extensive proteome remodeling with engagement of the autophagy pathway when proteasome capacity is limited.
- Our findings (i) indicate that thiol depletion and disruption of proteostasis through juglone’s dual redox cycling and alkylation activities are central to its phytotoxicity, (ii) cast doubt on previous reports that juglone targets a specific enzyme in plants or other organisms, and (iii) provide insight into how the chemical properties of allelopathic quinones shape their ecological roles.

## INTRODUCTION

Juglone (5-hydroxy-1,4-naphthoquinone; Fig 1a) is the phytotoxic compound produced by the black walnut tree (*Juglans nigra)*. It has a wide range of reported biological activities across plants, animals, fungi, and bacteria (Islam & Widhalm, 2020). Since it was proposed as the toxic constituent of black walnut a century ago (Massey, 1925), juglone has served as a model allelochemical for studying plant-plant interactions and has attracted significant interest for its agrichemical and medicinal properties (Widhalm & Rhodes, 2016).

**Fig. 1.**
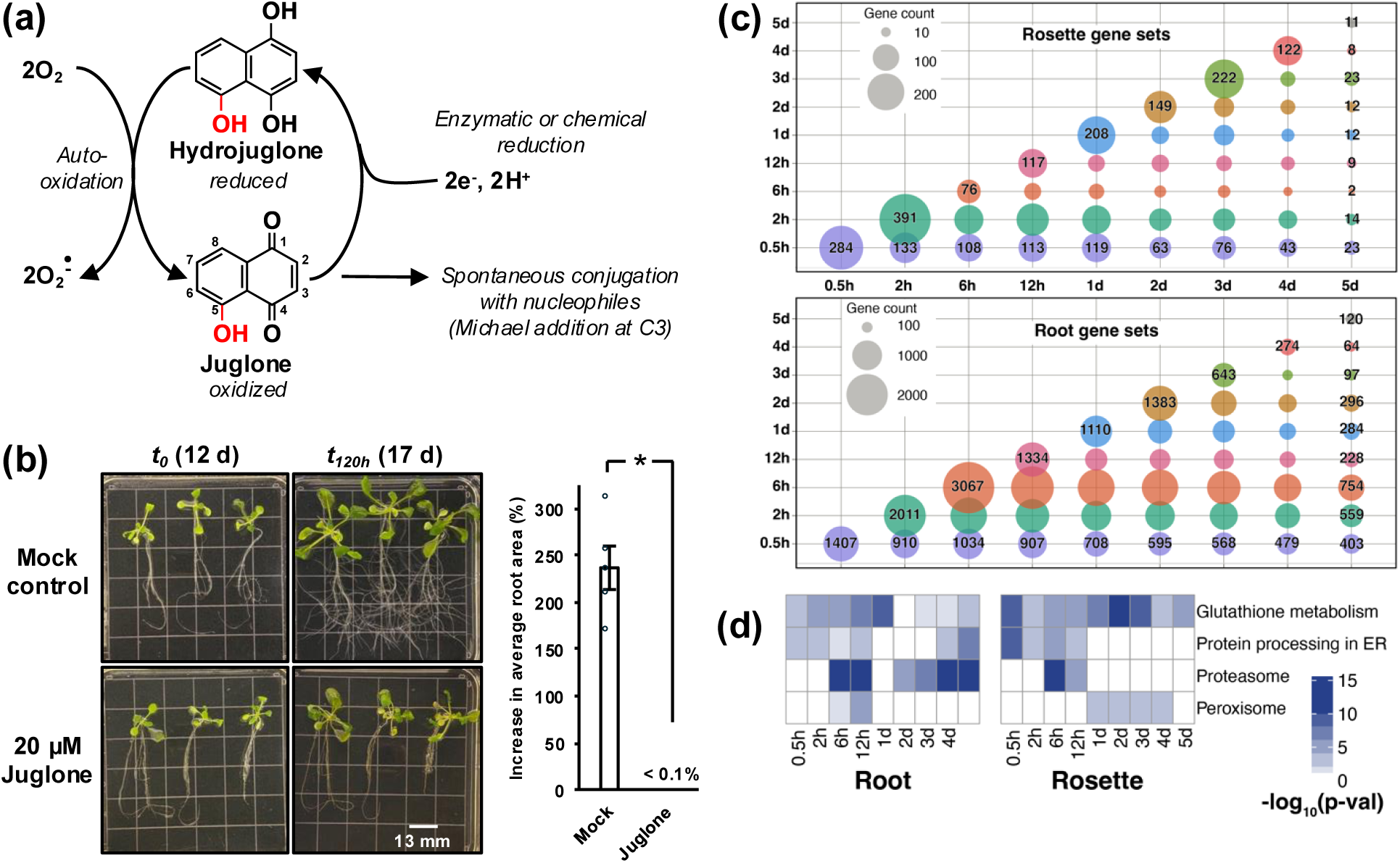
Juglone is a phytotoxic 1,4-naphthoquinone. (a) Predicted general mechanisms of action of juglone based on known quinone chemistry and biochemistry. Upon uptake, juglone undergoes redox cycling via reduction and subsequent re-oxidation by molecular oxygen, leading to the formation of reactive oxygen species. The hydroxyl group at C5 enhances the electrophilicity of C3, making juglone especially reactive with nucleophiles via Michael addition. (b) Juglone impairs root growth and foraging in Arabidopsis. Representative images show 12-d-old Arabidopsis transferred to ½ MS media containing 0 µM (mock) or 20 µM juglone at the start of treatment (*t_0_*) and 5 d after transfer (t_120h_). Grid squares are 13x13 mm. Quantification of the percent increase in average root area from t_120h_ relative to *t_0_* is shown to the right. Values are mean ± SE (n=5 with 3 plants per replicate); *p < 0.001, Student’s *t*-test. (c) Bubble plots showing the overlap of differentially expressed genes across time points in juglone-treated Arabidopsis rosettes and roots compared to organs from untreated plants sampled at *t_0_* (before transferring plants to treatment media). DEGs in response to juglone treatment remain persistent in the roots over the 5 days assayed, while the early (0.5-12 h) DEGs in the rosette return to their regular expression patterns after 1 d, indicating a recovery from the initial juglone shock. (d) Glutathione metabolism and proteasome pathways are upregulated throughout the 5-d treatment period in the roots. However, the rosettes switch to peroxisome-driven redox mitigation after 1-d post-juglone treatment.

In land plants, juglone is biochemically implicated as an inhibitor of plasma membrane H⁺-ATPase (Jose & Gillespie, 1998; Dayan & Duke, 2014). Juglone belongs to a class of compounds called the 1,4-naphthoquinones, which exhibit potent biological activities through their general ability to redox cycle and their electrophilic reactivity with nucleophiles (O’Brien, 1991; Klotz *et al*., 2014; Bolton & Dunlap, 2017) (Fig. 1a). The broad chemical reactivity of juglone with many molecular targets is reported across eukaryotes and prokaryotes. For example, juglone induces oxidative stress by depleting pools of plastoquinol and other prenyllipid antioxidants in *Chlamydomonas reinhardtii* (Nowicka *et al*., 2017, 2023). Juglone was also shown to disrupt electron transfer reactions essential for microbial nitrification through interference with hydroxylamine oxidoreductase (Akutsu *et al*., 2023). Moreover, juglone can irreversibly inhibit peptidyl-prolyl isomerases of the parvulin family, including Pin1 from *Escherichia coli*, *Saccharomyces cerevisiae*, *Malus domestica* (apple), and *Homo sapiens* (human) (Hennig *et al*., 1998; Chao *et al*., 2001; Yao *et al*., 2001). Similar results were found with urease from *Canavalia ensiformis* (jack bean) (Kot *et al*., 2010). Studies in human keratinocytes and *Rattus norvegicus* (rat) hepatocytes have additionally revealed juglone effectively depletes intracellular levels of reduced glutathione (GSH) (Ollinger & Brunmark, 1991; Inbaraj & Chignell, 2004) through oxidation of GSH to glutathione disulfide (GSSG), Michael addition of the L-cysteine thiol group in GSH to juglone to form a juglone-SG conjugate, or both.

Given juglone’s broad reactivity with molecular targets in species across the tree of life, we hypothesized that the phytotoxicity of juglone in plants arises not from inhibiting a specific protein or process, but rather from a more generalized disruption of cellular biochemistry. To investigate this, we combined comparative transcriptomics with biochemical tests, confocal microscopy, and quantitative proteomics to examine early and late responses of *Arabidopsis thaliana* to juglone. The findings from our study reveal a broader mechanism of juglone phytotoxicity more consistent with the general reactivity of 1,4-naphthoquinones and provide insight into how plants perceive and respond to redox active and electrophilic natural products in their environment.

## MATERIALS AND METHODS

### Plant growth

*Arabidopsis thaliana* (Col-0) seeds were sterilized in a glass desiccator with chlorine gas produced from 30 mL of household bleach and 5 mL of 12.1 N HCl for one hour. Sterilized seeds were plated on solid half-strength Murashige & Skoog (MS) medium containing 1% (w/v) sucrose, pH 5.7, and stratified at 4°C for at least two days. After stratification, seedlings, three per plate, were grown for 12 days under 16 hours of light and 8 hours of darkness in a growth room at a constant temperature of 22°C.

### Phenotyping root growth

Top-down photographs of individual plates with three Arabidopsis seedlings after treatments were taken in an imaging chamber equally lit from all sides. Plant images were processed in FIJI using a seven step imaging processing pipeline: 1) raw images were converted to JPG format, 2) image was duplicated, 3) duplicated image cropped to contain only the bottom 2/3 of the plate containing roots, 4) images were split by color channel into red, green, and blue images, 5) auto contrast threshold was applied to the blue image to remove dark non-background elements, 6) a mask of remaining non-background elements was created, 7) the percentage area covered by the mask, which appears as white on black background was measured. Cropping was identical for all photos taken on the same day. Any remaining leaf tissue in the cropped image was removed by the contrast threshold applied to the blue color channelsince it is significantly darker than root tissue in the blue channel. Each image’s percentage area represented the total sum of the root area from all three of the seedlings on each plate. Percent area data was processed by subtracting the percent area on day 0 from day 5, on a plate-by-plate basis for each plate’s percent root growth. Standard error for percent root growth was calculated for each treatment and a Tukey’s Multiple Comparison test with an Alpha=0.05 was used to contrast the mean root growth between groups.

### Tissue collection and RNA extraction for RNA-seq

12-day old Arabidopsis plants were transferred to solid half-strength MS media containing 1% sucrose dosed with 0 or 20 µM juglone. Juglone stock used to dose media was dissolved in acetonitrile and added to cooled media after autoclaving. Three plants were transferred to square petri dishes and placed back into growth room chamber until time of harvest. Rosettes were separated from roots at time of harvest and nine plants were pooled for each replicate. Tissue was flash frozen in liquid nitrogen and stored at -80°C until homogenization. Total RNA was extracted from 100 mg of homogenate with an RNeasy Plant Mini Kit (QIAGEN; Hilden, Germany) followed by an RNA Clean & Concentrator (Zymo Research Inc., Irvin, CA, USA). RNA purity and quantity were checked on the Thermo Scientific Nanodrop 2000c.

Library construction from 1 μg RNA, Illumina sequencing, and analyses of DEGs were performed by Novogene Corporation Inc. (Sacramento, CA, USA). Paired-end clean reads were mapped to the *Arabidopsis thaliana* release 44 version reference genome using HISAT2 software. For each sequenced library, read counts were adjusted using FPKM (Trapnell *et al*., 2010) and DEG analysis was performed using DESeq2 (Anders & Huber, 2010) with p-value adjusted using an FDR calculated with Benjamini–Hochberg (BH) methods (Benjamini & Hochberg, 1995). Genes were considered significantly differentially expressed if they had a BH-adjusted p-value of 0.005 and a log2 fold change of 1. Kyoto Encyclopedia of Genes and Genomes (KEGG) pathway enrichment analyses of DEGs was implemented by the clusterProfiler R package (Yu *et al*., 2012) and KEGG pathways with BH-adjusted p-value <0.05 were considered significantly enriched. The raw data will be submitted to the Sequence Read Archive and are available at the NCBI Sequence Read Archive.

### qPCR

The extracted RNA was subsequently treated with DNase I and purified using the *RNA Clean & Concentrator™ Kit* (Zymo Research Inc., Irvine, CA, USA), which includes an on-column DNase I digestion step, according to the manufacturer’s protocol. RNA integrity was assessed by agarose gel electrophoresis prior to downstream applications. Complementary DNA (cDNA) was synthesized using 500 ng of purified total RNA using the *5X All-In-One RT MasterMix* (Applied Biological Materials Inc., Richmond, BC, Canada), following the manufacturer’s guidelines. The cDNAs were further diluted 10-fold before being used for RT-qPCR. The NO APICAL MERISTEM/ARABIDOPSIS TRANSCRIPTION ACTIVATION FACTOR1/CUP-SHAPED COTYLEDONS2 (NAC) transcription factors *NAC53* (At3G10500) and *NAC78* (At5G04410) primers (Gladman et. al., 2016) were used to check expression relative to *ACTIN2* (At1G18780).

### LC-MS detection of juglone-SG conjugate formation in vitro

Juglone-GSH conjugate formation was analyzed using an Agilent 1260 high-performance liquid chromatography (LC) system coupled to an Agilent 1260 Infinity II diode array detector (DAD) and an Agilent 6120 single-quadrupole mass spectrometer. Separation was achieved on an Agilent Zorbax SB-C18 column (4.6 × 250 mm) maintained at 40°C. The mobile phase consisted of solvent A: 50 mM ammonium acetate with 0.1% formic acid in water, and solvent B: 50 mM ammonium acetate with 0.1% formic acid in methanol. The elution gradient was as follows: 0–12 min, 80% A; 12–51 min, linear gradient to 20% A; 51–60 min, linear gradient to 0% A; 60–66 min, return to 80% A; 66–75 min, 80% A, at a flow rate of 0.25 mL min^-1^.

UV absorbance was monitored at 260 nm. Authentic standards of GSH and juglone were used to determine their retention times of 9.6 min and 49.8 min, respectively. A new peak corresponding to the putative juglone-SG conjugate was observed at 29.8 min. Mass spectrometry detection was conducted using electrospray ionization in negative ion mode with the following source conditions: gas temperature 350°C, drying gas flow 12 L min^-1^, nebulizer pressure 35 psig, and capillary voltage 3000 V. Mass spectra were acquired in full scan mode from *m/z* 300–600 using a fragmentor voltage of 50 V. Ion extraction was performed for *m/z* 478 (expected [M-H]⁻ of the GSH-juglone conjugate) and *m/z* 306 (GSH). Data were analyzed using the Chemstation software suite.

### Extraction and detection of juglone-SG in planta

Seedlings were grown for 12 d and transferred to treatment plates containing 20 µM juglone as described above. Samples were subsequently incubated for 0, 2, and 6 h. Following treatment, roots were harvested, flash-frozen, and processed. A total of 100 mg pulverized root tissue was extracted in 0.6 mL methanol:water (50:50, v/v) for 1 h, and 20 µL of the resulting extract was used for analysis.

Juglone-SG conjugate analyses were performed using an Agilent LCMS-6460 triple quadrupole mass spectrometer equipped with an Agilent 1200 Infinity HPLC system and Zorbax reversed-phase C18 column (2.1 × 50 mm, 3.5 μm). The chromatographic separation was achieved using a mobile phase of water containing 0.1% formic acid (solvent A) and acetonitrile containing 0.1% formic acid (solvent B). The following linear gradient elution profile was used: 0 min 5% of B, linear gradient increase to 100% of B over 6 mins, followed by a 0.5-min hold at 100% B. The column was subsequently re-equilibrated with 5% B for 3 min between runs. A flow rate of 0.3 mL min^-1^ was maintained throughout the analysis, with the column temperature set at 30°C. The injection volume for all the samples was 20 μL.

Juglone, GSH, and the in-vitro synthesized juglone-SG conjugate were analyzed in both positive and negative electrospray ionization (ESI) modes. The negative ionization mode yielded the highest abundance for the respective analytes: juglone (*m/z* 173), GSH (*m/z* 306), and juglone-SG conjugate (*m/z* 478). The MS/MS fragmentation of the conjugate in negative ion mode produced characteristic transitions of *m/z* 478→202, 478→205, and 478→272. The Multiple Reaction Monitoring (MRM) was set to monitor these transitions, with the following conditions: electrospray source with a nitrogen gas temperature of 325°C (flow rate 7 L min^-1^), sheath gas temperature of 250°C (flow rate 7 L min^-1^), capillary voltage of 3800 V, nozzle voltage of 500 V, and fragmentor voltage of 80 V. Nitrogen served as the collision-induced dissociation (CID) gas, with collision energies set at 10 eV for transitions to *m/z* 202 and 272, and 20 eV for transition to *m/z* 205.

### MALDI analysis

BSA from Sigma (A7517) was dissolved in 50mM potassium phosphate buffer pH 7.7 and 2 mM EDTA at a concentration of 1.25 mg mL^-1^. Juglone dissolved in MeOH was added to the buffer for a final concentration of 50 µM. Juglone and BSA were incubated in the dark for 40 minutes at room temp before juglone was removed with a GE Healthcare Shepadex PD-10 column and 0.5 ml fractions were collected. Fractions with highest protein content were pooled and diluted to 1 mg/mL then flash frozen in liquid nitrogen and stored at -80° C until analysis. Protein samples were diluted 1:100 in water with 0.1% trifluoroacetic acid. Sinapinic acid (Sigma Aldrich) was used as the matrix (10 mg/mL in 50:50 acetonitrile:water with 0.1% TFA). Equal parts sample and matrix were mixed and applied to a 384-well stainless-steel plate via dried droplet application. The samples were then dried *in vacuo*. Samples were loaded into a 4800 MALDI TOF/TOF Analyzer (Applied Biosystems/MDS SCIEX) with 4000 Series Explorer v3.5 software for acquisition. The instrument was calibrated prior to sample spectrum collection. The samples were acquired automatically in MS linear positive ion mode, with a fixed laser intensity of 7900, in a range of 30,000-150,000 Da.

### Monochlorobimane staining and imaging

Twelve-day old Arabidopsis plants were transferred to solid ½ MS 1% sucrose dosed with juglone, lawsone, or mock. Quinone stocks were dissolved in 100% methanol. Arabidopsis plants were incubated on quinone or mock treated plates for 2 h before transferring to PBS (136.9 mM NaCl, 268.3 µM KCl, 10.1 mM Na_2_HPO_4_, and 1.8 mM KH_2_PO_4_) pH 7.4 buffer with 2.8 µg mL^-1^ monochlorobimane. Plants were stained for 1 h in the dark before they were rinsed in PBS buffer and imaged. Photos were taken on Zeiss LSM 880 Upright Confocal microscope with 405 nm excitation and 491 nm emission wavelengths. Relative fluorescence was quantified by uploading the CZI files to FIJI and quantifying the average pixel intensity of the first 300 µm of the root tip and subtracted out background intensity.

### DTNB assays

Arabidopsis plants were grown for 12 days in ½ MS media and 1% sucrose. Roots were separated from rosette and flash frozen in liquid nitrogen. Approximately 200 mg of the root tissue was ground in 2 mL of 100 mM Tris-HCl, pH 8.3, 10% (v/v) glycerol with protease inhibitor minitablets from Thermo (product number: 88665) in a dunce homogenizer for 20 minutes. Homogenate was centrifuged for 10000 x g for 10 minutes at 4°C and the supernatant was desalted in a GE Healthcare Shepadex PD-10 Column into 50 mM potassium phosphate buffer pH 7.7. Protein was quantified using the Quick Start Bradford protein assay kit and BSA (Bio-Rad). 4.5 µg of protein was incubated in potassium phosphate buffer with 0, 12.5, or 50 µM of juglone or lawsone for 45 minutes in a final volume of 50 µL in the dark at room temp. After incubating, 50 µL of 1 mM DTNB in phosphate buffer was added and pipetted to mix. Absorbance was measured at 412 nm on nanodrop in a quartz cuvette after 25 minutes of incubating with DTNB reagent. For assays with BSA, the protein was incubated with juglone, lawsone, or mock as described for MALDI analysis. Desalted protein was diluted to 1 mg mL^-1^ protein. Exactly 37.5 µL of 6.3 mM DTNB reagent was then added to 750 µL of the protein and pipetted to mix before recording absorbance at 412 nm every 10 seconds over 5 minutes in a quartz cuvette.

### PAL assays

Arabidopsis plants were grown for 12 days in ½ MS media and 1% sucrose. Roots were separated from rosette and flash frozen in liquid nitrogen. Approximately 200 mg of the root tissue was ground in two ml of 100 mM Tris-HCl, pH 8.3, 10% (v/v) glycerol with protease inhibitor minitablets from Thermo Scientific (catalog number 88665) in a dunce homogenizer for 20 minutes. Homogenate was centrifuged for 10000 x g for 10 minutes at 4°C and the supernatant was desalted in a GE Healthcare Shepadex PD-10 Column into Tris-HCl buffer without protease inhibitor. Protein was quantified using the Quick Start Bradford Protein assay kit and BSA. 15 µg of root protein extracts were incubated in a 485 µl reaction volume with 200, 150, 100, 75, 50, 25, 10, 5 and 0 µM juglone in Tris-HCL buffer for 45 minutes at 37°C. Assays were initiated with the addition of 15 µL of 133.33 mM Phe in Tris-HCL buffer. Enzyme assays were incubated at 37°C in the dark for 45 minutes. Assays were stopped with the addition of 50 µL of glacial acetic acid and metabolites were extracted in 750 µL of ethyl acetate. 500 µL of the ethyl acetate fraction was evaporated and redissolved in 50% MeOH. Samples were briefly spun down at 15,000 x g. Product *trans*-cinnamic acid, was analyzed on Agilent Technologies 1260 Infinity HPLC-DAD at wavelength 273 nm. Metabolites were separated on a ZORBAX SB-C18 StableBond Analytical 5-Micron column with a flow rate of 1 mL/min with solvent A (0.1% FA in water) and solvent B (0.1% FA in acetonitrile). The separation method was initiated by 90% A for 2 minutes before it was decreased to 75% after 8 minutes, and then 45% after 22 minutes. Mobile phase A was held at 10% from 24-25 minutes before it was increased back to 90% over 2 minutes and held for 8 minutes.

### Protein extraction for proteomics

Whole 12-d-old Arabidopsis plants were ground using Research Products International Corp Micro-Pestles and liquid nitrogen before 100 mg was aliquoted into LoBind Eppendorf tubes. One mL of extraction buffer (0.5 M HEPES (pH 7.5), 5 mM Na_2_EDTA, 2 mM dithiothreitol (DTT), and 1x protease inhibitor) was added to the tissue and further ground with a dunce homogenizer under ice-cold conditions. Samples were incubated for 20 min in ice. Subsequently, samples were centrifuged at 16,000 x *g* for 20 min at 4 C. The supernatant was separated, and proteins were precipitated using a 80% (v/v) acetone at -20°C. Next, the protein pellets were washed with 400 µl of 100% acetone before spinning at max speed for 20 min at 4°C. The remaining protein pellets were dried down in a vacuum and then dissolved in 100 µL of 25 mM ammonium bicarbonate buffer (AMBIC buffer) supplemented with 8 M urea. The proteins were extracted as described in (Thirumalaikumar *et al*., 2023). Briefly, 100 ug of protein was subjected to digestion. Six µL of 200 mM DTT was used for the reduction step. 24 µL of 200 mM iodoacetamide in the dark was used for the alkylation step. Residual IAA was quenched with DTT treatment. Protein extracts were diluted with 700 µL of AB lysis solution and 5 µL of Trypsin/Lys-C protease (1:100) were added and incubated overnight at 37°C. Another 2.5 µL of Trypsin/Lys-C was used to top up the digestion for 5 hours. Trypsin was stopped using trifluoracetic acid (TFA). Samples were dried down under a vacuum at RT before resuspending the pellet in the tube with 100 µl of AMBIC buffer. Peptides were desalted using Pierce C18 Spin Tips & Columns following the manufacturer’s instructions. Eluted samples were vacuum dried at 45°C, and the pellet was dissolved in 20 µl of 5% acetonitrile. Peptide concentration was quantified using the Pierce Quantitative Colorimetric Peptide Assay Kit adjusted to 1 µg µL^-1^.

### Data-dependent proteomics analysis

The LC method and the MS settings were kept as same as in (Grantz *et al*., 2024). Peptides were analyzed by a Orbitrap Lumos (Thermo Fisher Inc., Waltham, MA, USA) coupled with an Ultimate 3000 HPLC (Thermo Fisher Inc., Waltham, MA) and an Aurora Ultimate analytical column (IonOpticks, Victoria, AUS). Approximately 0.6 μg of peptides were loaded onto a trap column (C18) and separated on a 25 cm nanoflow UHPLC IonOpticks C18 column. A 130-minute elution gradient was constructed by mixing mobile phase solvent A (0.1% FA in water) with solvent B (80% ACN, 0.1% FA in water). The gradient started with 2% solvent B, increasing to 12% at 1.6 min, 25% at 80 min, 35% at 100 min, 45% at 105 min, and 95% at 120 min. The gradient was held at 95% for 5 min before reverting to 2% at 125.1 min, and 5% at 130 min. All data were acquired using the Lumos Orbitrap MS (Thermo Fisher Scientific). The Orbitrap mass analyzer collected data using a higher-energy collisional dissociation fragmentation scheme. For MS scans, the scan range was from 375 to 1500 *m/z* at a resolution of 120,000, with an automatic gain control (AGC) target set at 4 × 10^5^, a maximum injection time of 50 ms, dynamic exclusion of 60 s, and an intensity threshold of 5.0 × 10^3^. Data were acquired in data-dependent (DDA) mode with a cycle time of 3 s per scan. MS2 data were collected at a resolution of 7,500. The integrity and performance of the mass spectrometer were monitored using HeLa standards before and after the experimental runs.

### Data-independent acquisition (DIA) proteomics analysis

Peptides were analyzed using Brucker trapped ion mobility-time-of-flight high-throughput (timsTOF HT) (Bruker Daltonics Gmbh) MS coupled with the nano Elute 2 (Bruker Daltonics GmbH) reverse phase liquid chromatography system coupled to a Captive Spray 2 ion source. Briefly, 100 ng of peptide from the Arabidopsis root and seeding samples were analyzed using the nano Elute 2 LC system using a 60-min active gradient. Peptides were separated using a 2-column (trap and analytical) separation technique. Peptides were first loaded onto the trap cartridge (5mm x 300 µM, 5 µm particle size, and 100 Å pore size, ThermoFisher Scientific) and subsequently into reprosilC18 (25 cm × 150 µM, 1.9 µm particle size, and 120 Å pore size; Brucker Daltonics Gmbh, Part No:1895619) analytical column. Peptides were then eluted over a 70-min gradient ranging from 0-26% Solvent B (0.1% FA in ACN) for 50 min, from 26-32% Solvent B (0.1% FA in ACN) for 10 min, and set to 95% B for another 10 min to wash. The analytical column was equilibrated for the starting condition for the next 10 min, at a flow rate of 350 nL min-^1^. The column temperature was maintained at 50°C. Peptides were sprayed into the MS using a 20 µm PepSep emitter (Bruker Daltonics GmbH). All data were acquired under the DIA-PASEF (data independent acquisition-Parallel Accumulation Serial Fragmentation) mode (Meier *et al*., 2020). To minimize carryover and ensure consistent column conditions, sample injections were alternated with blank injections (solvent A). The mass spectrometer’s performance was monitored daily using a HeLa protein standard. Before sample analysis, the instrument was calibrated for both mass and ion mobility using three reference ions from the Agilent ESI-L Tuning Mix (*m/z* 622, 922, and 1,222).

### Data-dependent acquisition (DDA) proteomics analysis

Data-dependent acquisition (DDA) data were analyzed using Proteome Discoverer, and DIA data were analyzed using Spectronot following the default settings. Briefly, for the DDA data, an in-house list of common contaminants was added to the search. Sequest HT tool was used to assign the peptides, allowing a maximum of two missed tryptic cleavages, a minimum peptide length of six, a precursor mass tolerance of 10 ppm, and a fragment mass tolerance of 0.02 Da. Carbamidomethylation of cysteines and oxidation of methionine were specified as static and dynamic modifications, respectively. A false discovery rate of high confidence validated peptide spectral matches was used for the downstream analysis.

Label-free quantification based on MS1 precursor ion intensity was performed in Proteome Discoverer. The normalized protein abundances were calculated among the measured samples, and values were log2-transformed and imputed following a normal distribution pattern. For the DIA data we have used the spectronaut default settings, and the protein quantification data were exported. R-studio, Perseus, and GraphPad were used to generate the graphs and compute the data. Adobe Illustrator and Microsoft PowerPoint were used to present graphical visuals.

## RESULTS

### Juglone activates stress pathways consistent with responses to its redox chemistry and electrophilicity

Despite juglone’s widely reported phytotoxic effects (Islam & Widhalm, 2020), understanding of the stress incurred by plants exposed to juglone and the subsequent responses remains limited. A previous study reported expression profiles for rice seedling roots (*Oryza sativa* L. cv. TN-67) after being exposed to juglone for 1 and 3 h (Chi *et al*., 2011). However, plants growing near walnut trees often must manage juglone exposure for sustained periods of time as the allelochemical is continuously leached into the soil from roots and falling fruits. Therefore, to better understand how plants initially and continually respond to juglone, we captured early-to-late transcriptome responses in separated rosettes and roots of 12-d-old Arabidopsis (Col-0) after 0, 0.5, 2, 6, 12, 24, 48, 72, 96, and 120 h following transfer to media containing 20 µM juglone (Supporting Information Fig. S1). This concentration is orders of magnitude lower than the 0.25-3.25 mM range reported in the soil under black walnut trees (De Scisciolo *et al*., 1990), but was sufficient to arrest growth, inhibit root foraging, and induce visible signs of injury without killing the plant (Fig. 1b), thus allowing for the extraction of suitable quality RNA for RNA-seq.

Using comparisons of gene expression in roots or rosettes from plants treated with juglone relative to gene expression immediately before juglone exposure (*t_0_*), we analyzed differentially expressed genes (DEGs) across early (0.5, 2, 6, 12, and 24 h) and late (24, 48, 72, 96, and 120 h) response time points. Our analysis revealed that in rosettes, 2350 DEGs were common across the early time points compared to *t_0_*, while 3772 DEGs were shared in the late responses (Fig. S2). Roots exhibited a similar transcriptional response, with 1774 DEGs common among early time points compared to *t_0_*and 5021 DEGs persisting throughout the late phase (Fig. S2). Tracking the persistence of DEGs across the time points shows that in roots the genes responding to juglone early (within 6 h) largely remain DE at later time points (Fig. 1c) while the early DEGs in the rosettes are largely not DE after 24 hours (Fig. 1c).

Kyoto Encyclopedia of Genes and Genomes (KEGG) pathway enrichment analysis of DEGs highlighted an immediate molecular response that matches juglone’s propensity to redox cycle and react with nucleophiles. In both roots and rosettes, the categories “glutathione metabolism,” “protein processing in the ER,” and “proteasome” were significantly enriched among the upregulated DEGs within the first 6 h of juglone treatment compared to *t_0_* (Figs. S3,4). Both “glutathione metabolism” and “proteasome” remained significantly enriched among the upregulated DEGs by 24 h in both roots and rosettes compared to *t_0_*. From 0.5-12 h, the category “ascorbate and alderate metabolism” was significantly enriched among the upregulated genes in roots relative to *t_0_*(Fig. S3). Categories for “cysteine and methionine metabolism” and “core-autophagy (ATG) genes” were also enriched at various times in the early time points.

Across the later sampled time points, 24-120 h, “proteasome” remained enriched among the upregulated DEGs in roots, while “glutathione metabolism” remained enriched in rosettes compared to *t_0_*. The proteasome pathway in KEGG includes the set of genes involved in rapid degradation of ubiquitinated proteins. In the rosettes, in the later time points the proteasome pathway enrichment is replaced by the peroxisome (Fig. 1d), e.g., by the upregulation of acyl-CoA oxidases (ACX1, ACX2, ACX3 and ACX4) that catalyze the first step in β-oxidation of fatty acids, which eventually increases the supply of NADPH, an essential cofactor for regenerating GSH. Only the category “glucosinolate biosynthesis” was enriched among the downregulated genes in roots across early and late time points compared to *t_0_*. No other remarkable discernible patterns were observed among the enriched categories from the downregulated genes in roots or rosettes (Figs. S5-6). We also analyzed DEGs across early and late time points in juglone-treated versus mock-treated roots and rosettes sampled at the same time following treatment (Figs. S7-11), which revealed similar trends of enriched categories as those comparisons made relative to *t_0_*(Figs. S2-6).

### Juglone disrupts thiol homeostasis in glutathione (GSH) and proteins

Based on the enrichment of glutathione, cysteine, methionine, and sulfur metabolism pathways after juglone exposure, we hypothesized that juglone perturbs thiol homeostasis in peptides and proteins *in planta*. We first reassessed juglone’s reactivity in vitro with the antioxidant tripeptide GSH (𝛄-glutamyl-cysteinylglycine) using 5,5’-dithiobis-(2-nitrobenzoic acid) (DTNB), which reports concentrations of reduced sulfhydryl groups. Incubation of GSH with juglone resulted in a stoichiometric decrease in detectable thiols, whereas incubation with lawsone (2-hydroxy-1,4-naphthoquinone), the principal bioactive quinone responsible for the orange-dye properties of *Lawsonia inermis* (henna), did not differ from the mock (Fig. S12a). While lawsone can undergo redox cycling like juglone (Osman & van Noort, 2003; Sauriasari *et al*., 2007), it cannot spontaneously conjugate with GSH because its C3 position is saturated in its favored “keto” form (Klaus *et al*., 2010) (Fig. S12b). Thus, our results support previous work comparing the chemistries of juglone and lawsone (Doherty *et al*., 1987; Kot *et al*., 2010) and work reporting that juglone cytotoxicity involves nucleophilic addition of juglone to GSH to form a juglone-SG conjugate (Ollinger *et al*., 1989; Inbaraj & Chignell, 2004). To further validate the formation of juglone-SG, we analyzed the product of the in vitro juglone and GSH reaction by high-performance liquid chromatography coupled with diode-array detection and single-quadrupole mass spectrometry (HPLC-DAD-MS). A novel compound with a mass of *m/z* 478 was detected, consistent with the expected mass of juglone-SG (Fig. S12c-e). It is important to note that conjugation occurred in the absence of glutathione-*S*-transferases (GSTs) or any other added enzyme.

Having verified that juglone spontaneously forms a GSH conjugate in vitro, we next asked whether a juglone-SG conjugate pool is detectable in roots exposed to juglone. To test this, we analyzed root extracts from Arabidopsis treated with juglone by LC-MS/MS. Root extracts from 12-d-old Arabidopsis plants exposed to juglone for 2 h or 6 h were analyzed by multiple-reaction-monitoring (MRM). The *m/z* 478 precursor ion produced an ion spectrum with a dominant fragmentation pattern of *m/z* 202, 205, and 272. The transition from *m/z* 478 to 205 was the most abundant and was used for measurement. The *m/z* 205 ion corresponds to a fragment in which intact juglone remains covalently attached to the sulfur atom of GSH (Fig. 2a). The *m/z* 272 ion is a diagnostic GSH fragment corresponding to deprotonated γ-glutamyl-dehydroalanyl-glycine (Dieckhaus *et al*., 2005; Xie *et al*., 2013; Huang *et al*., 2015). Quantification based on the *m/z* 478 to 205 transition revealed a pool of 8.3 pmol juglone-SG mg^-1^ FW in roots 2 h post-exposure and decreasing by approximately 25% after 6 h (Fig. 2b).

**Fig. 2.**
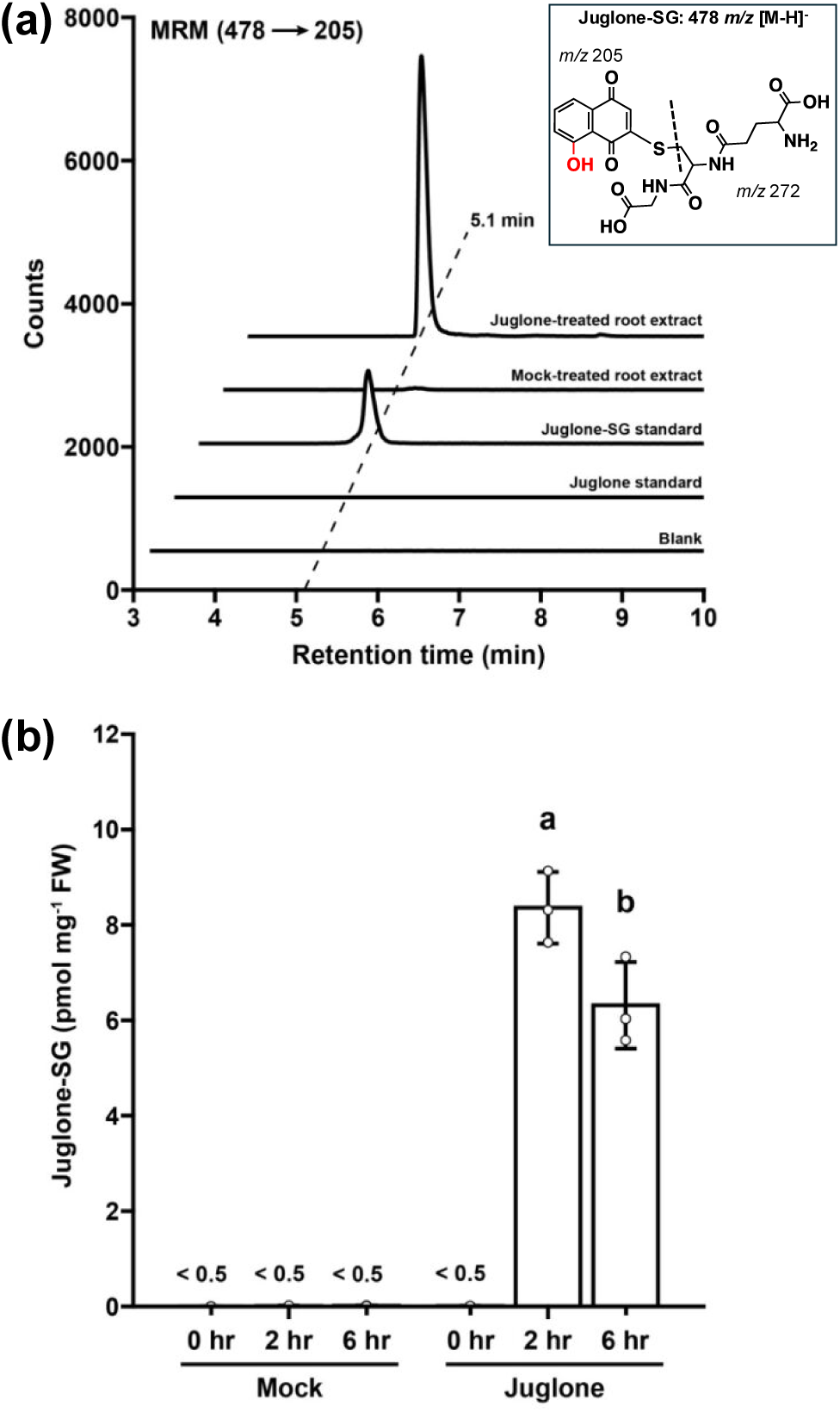
Detection and quantification of the juglone-glutathione (Jug-SG) conjugate in juglone-treated Arabidopsis roots. (a) Chromatograms of juglone, an in vitro synthesized juglone-SG conjugate, and extracts from Arabidopsis roots treated with juglone or mock treatments. Inset shows the structure of juglone-SG. The dashed line indicates fragmentation along the C-S bond of the GSH moiety in juglone-SG to produce the *m/z* 205 and 272 ions that correspond to the juglone thioether core and the deprotonated γ-glutamyl-dehydroalanyl-glycine fragment, respectively. (b) Quantification of juglone-SG from mock and juglone-treated Arabidopsis roots after 2 and 6 hrs. Bars show mean ± SE (n = 3); letters show statistical differences determined by a one-way ANOVA with Tukey’s post hoc analysis (p<0.05).

Next, to test whether juglone’s reactivity decreases pools of GSH in roots, we stained quinone-treated Arabidopsis roots with monochlorobimane (MCB), a fluorogenic probe for GSH. After 2 h of juglone exposure, MCB fluorescence was reduced by more than 90%, indicating depletion of free GSH compared to mock (Fig. 3a-d). In contrast, roots treated with lawsone retained MCB fluorescence comparable to mock controls. Given published estimates that total root GSH pools are in the millimolar range (e.g., Meyer et al., 2001), which is substantially larger than the pool of juglone-SG we detected in roots (Fig. 2b), the extent of MCB signal loss cannot be fully explained by measured juglone-SG. Either juglone-SG is rapidly catabolized, and/or it reflects GSH consumption via hydrojuglone redox cycling and associated thiol oxidation. Nonetheless, our findings demonstrate that juglone is taken up rapidly and significantly depletes the pool of GSH in Arabidopsis roots.

**Fig. 3.**
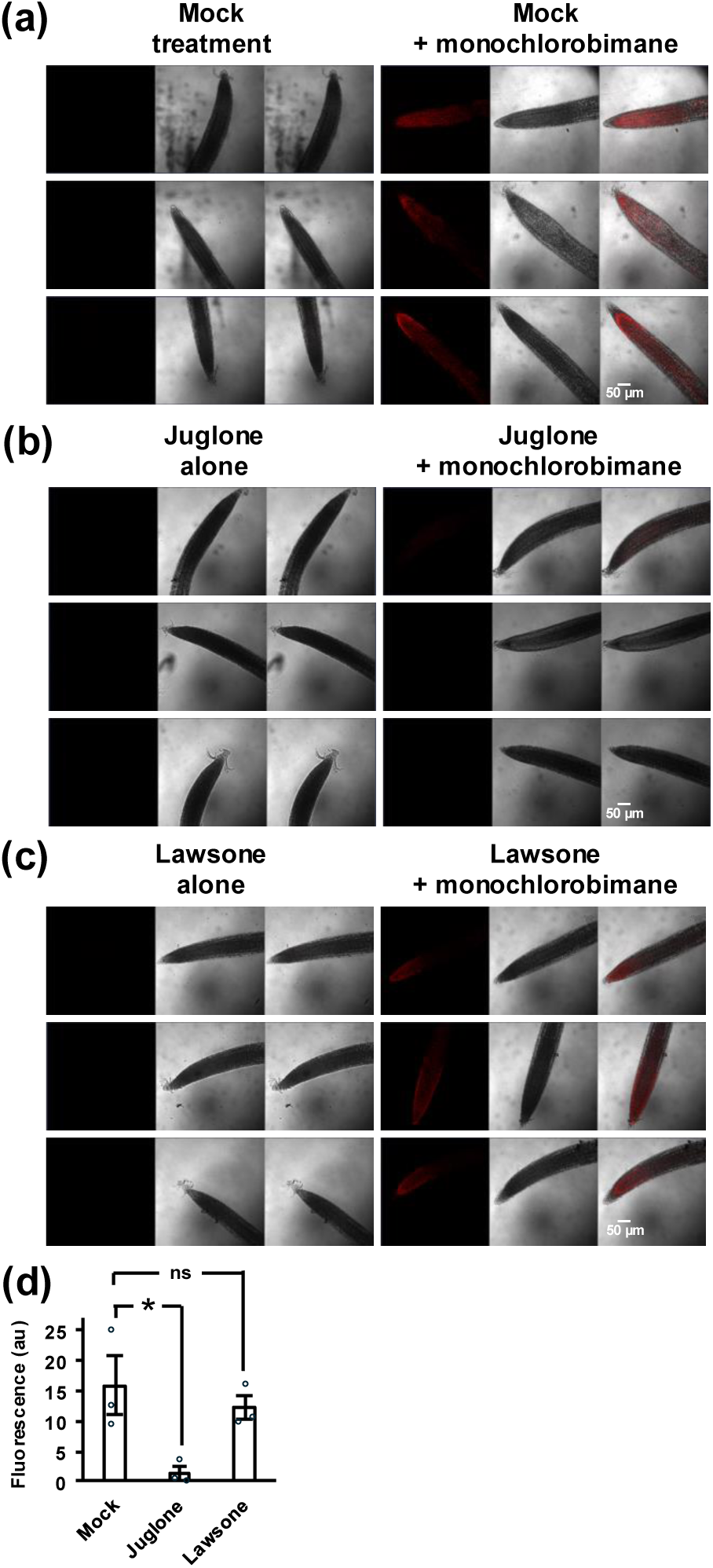
Juglone, but not lawsone, depletes free glutathione (GSH) pools in Arabidopsis roots. (a-c) confocal images show 12-d-old Arabidopsis roots treated with 20 µM quinone (juglone or lawsone) in ½ MS media ± 12.35 µM monochlorobimane (MCB) for 2 h. Fluorescence (red) indicates the presence of free GSH. Representative images of roots treated with mock (a), juglone (b), or lawsone (c), with and without MCB. Scale bars = 50 µm. Juglone treatment markedly reduced fluorescence, indicating depletion of free GSH in planta. (d) Quantification of fluorescence from MCB treatment. Bars show mean ± SE (n = 3); *p < 0.05, Student’s *t*-test; ns = not significant.

We next investigated whether juglone’s propensity to alkylate with thiol groups extends to proteins. We reasoned that if juglone can spontaneously conjugate with the cysteine thiol in the tripeptide GSH, then it should be able to covalently modify accessible cysteine residues in any protein. This idea is supported by previous work demonstrating that enzymes like urease (Kot *et al*., 2010) and peptidyl-prolyl isomerase (Hennig *et al*., 1998) are irreversibly inhibited by juglone. To investigate this further, we used bovine serum albumin (BSA) as a model to look for direct evidence of conjugation. BSA is a monomeric protein containing 35 cysteines that form 17 pairs of disulfide bonds, leaving only one cysteine residue, Cys-34, that exists as a free thiol group (Fig. S13a). Moreover, Cys-34 in BSA and other serum albumins is located on the protein surface and can be covalently modified by the electrophilic acrylamide-based fluorophore, acrylodan (Narazaki *et al*., 1997). We first tested thiol availability by quantifying reduced sulfhydryl groups using DTNB. A 1.25 mg mL^-1^ solution of BSA was prepared, pre-incubated with 50 µM juglone, lawsone, or mock treatment for 45 min, and desalted before quantifying thiol availability with DTNB. Whereas lawsone-treated BSA had an equal amount of unmodified thiol groups compared to the mock-treated control, the BSA pre-treated with juglone had almost no free sulfhydryls (Fig. S13b). To determine if BSA was covalently modified by juglone like GSH, we examined the masses of mock and juglone-treated samples using matrix-assisted laser desorption/ionization time-of-flight (MALDI-TOF) mass spectrometry. We detected a mass increase from 66,409 Da in the untreated mock to 66,583 Da in the juglone-treated BSA, consistent with alkylation by a single juglone molecule (174 Da) (Fig. 4a,b and Fig. S13c,d).

**Fig. 4.**
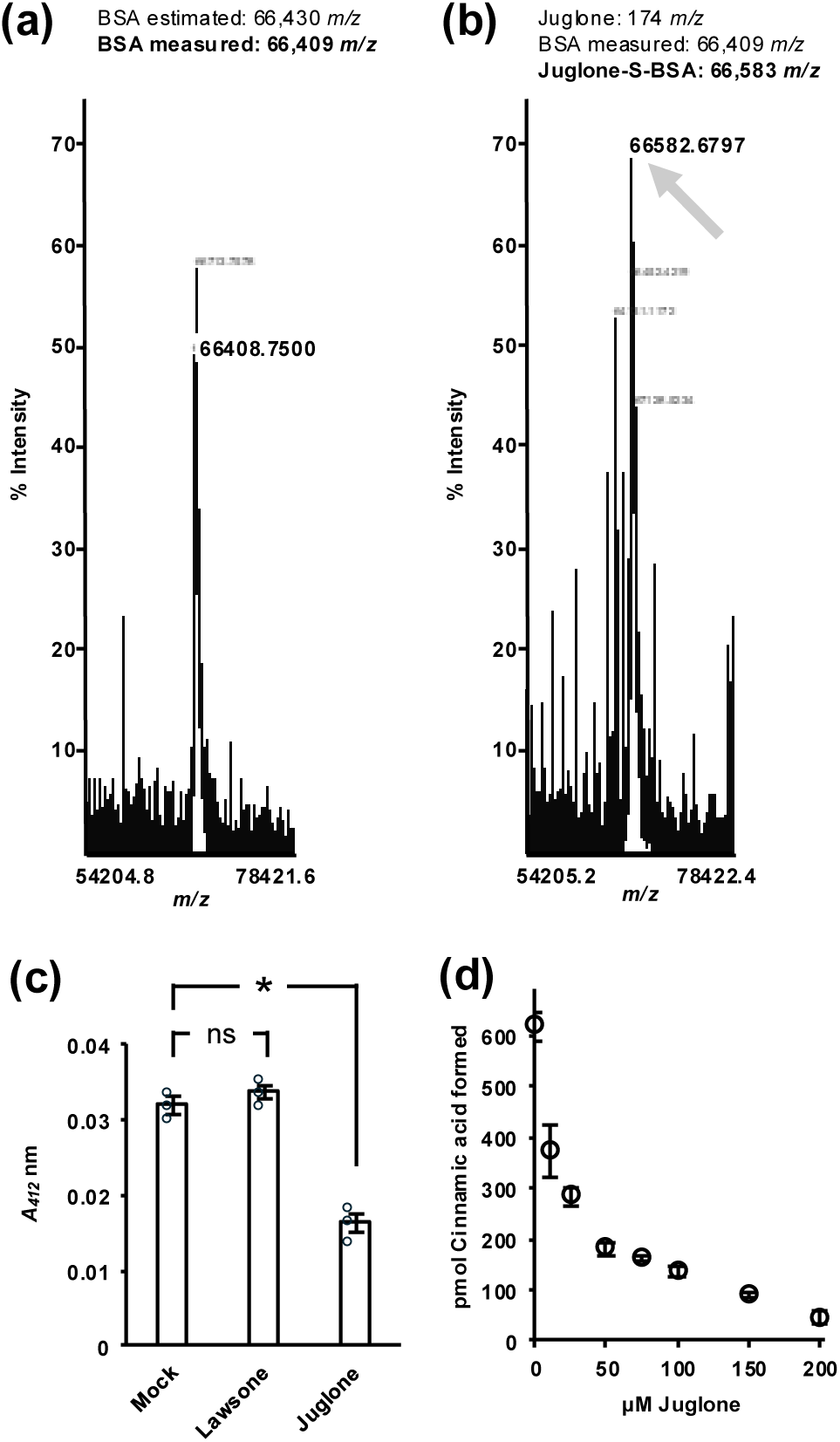
Juglone depletes available protein sulfhydryls and decreases the amount of phenylalanine ammonia-lyase (PAL) activity in root extracts. (a,b) MALDI-TOF mass spectrometry of BSA alone (a) or incubated with juglone (b). A new peak (arrow) corresponding to a mass increase of ∼174 Da in the juglone-treated sample is consistent with the covalent addition of one juglone molecule (174 *m/z*) to BSA (66,583 *m/z* vs. unmodified 66,409 *m/z*). Full spectra are presented in Supplementary Fig. S13. (c) Quantification of free thiol content in total protein extracts (4.48 mg mL^-1^) from 12-d-old Arabidopsis roots treated with 50 µM juglone, lawsone, or mock control for 5 min. Bars show mean ± SE (n = 3); *p < 0.05, Student’s *t*-test; ns = not significant. (d) In vitro PAL activity assay (45 min) using 12-d-old Arabidopsis root crude protein (15 µg) incubated with increasing concentrations of juglone. Juglone lowered the amount of PAL activity detected in a dose-dependent manner, as indicated by decreased formation of cinnamic acid from supplied phenylalanine. Data are mean ± SE (n = 3).

With evidence that juglone can covalently modify cysteine residues in both a peptide (GSH) and a protein (BSA), we next investigated whether juglone depletes total thiols in a complex protein sample. A desalted crude protein extract, which is comprised of myriad proteins, was prepared from roots of 12-d-old Arabidopsis and incubated with 50 µM juglone, lawsone, or mock treatment. Based on DTNB measurement of free sulfhydryls, the lawsone-treated crude root protein was found to have an equal amount of available thiols compared to the mock-treated control, while the crude protein treated with juglone had approximately half as many (Fig. 4c). This suggests that juglone, whether it be through covalent modification, oxidation, or a combination of both, can disrupt thiols in a complex protein sample. Next, we predicted that if juglone broadly modifies thiols, then any enzyme with free cysteines will potentially exhibit reduced activity in the presence of juglone. To test this, we assayed phenylalanine ammonia-lyase (PAL) activity in the same root protein extracts across a range of juglone concentrations. PAL has eight cysteine residues, most of which appear to be located on the surface of the protein (Fig. S14). Indeed, the rate of cinnamic acid formation decreased from 13.73 pmol min^-1^ at 0 µM juglone to 1.02 pmol min^-1^ at 200 µM juglone in a dose-dependent manner (Fig. 4d). Given that we were able to predict juglone would lower the amount of PAL activity based on the presence of free cysteine residues in PAL, this suggests juglone does not selectively inactivate any particular enzyme. Instead, juglone appears to broadly modify sulfhydryl groups in peptides and proteins via conjugation, oxidation, or both, causing thiol depletion and widespread loss of cellular enzyme activities.

### The proteasome stress regulon is induced in response to juglone

Considering the broad reactivity of juglone with cysteine residues and the enrichment of genes upregulated by juglone that are linked to the processing of misfolded proteins, the proteasome, and autophagy (Fig. 1d, Figs. S3,4), we hypothesized that juglone elicits proteotoxic stress in Arabidopsis. Responding to proteotoxicity involves activating and coordinating several mechanisms, including the 26S proteasome to degrade ubiquitinated misfolded proteins, protein chaperones to help with protein folding and minimizing aggregation, and autophagy to clear insoluble protein aggregates and recycle damaged cellular components (Raffeiner *et al*., 2023). In Arabidopsis, the response to proteotoxicity is controlled by the proteasome stress regulon (PSR), which is a network of genes regulated by the NAC transcription factors NAC53 and NAC78 (Gladman *et al*., 2016).

Examination of our juglone time series RNA-seq data revealed that *NAC53* transcripts accumulated predominantly in rosettes, whereas *NAC78* was more strongly induced in roots (Fig. 5a). NAC53 and NAC78 have previously been shown to form both homo- and heterodimers that bind shared cis-elements within the promoters of PSR genes (Gladman *et al*., 2016). We further validated changes in *NAC53* and *NAC78* expression by qPCR using RNA extracted from roots and rosettes of 12-d-old Arabidopsis at 2 h and 12 h following exposure to 20 µM juglone (Fig. S15a). Expression of *NAC53* increased 1.8- and 2.0-fold in roots after 2 and 12 h, respectively, though only the latter was statistically significant. Consistent with the RNA-seq data, in rosettes, *NAC53* expression was even more highly induced, increasing by 4.5- and 5.1-fold at 2 and 12 h, respectively. Meanwhile, expression of *NAC78* significantly increased 2 h and 12 h following juglone exposure. In rosettes, *NAC78* expression increased by 1.4-1.6-fold, while in roots, expression increased 2.0-fold after 2 h and 4.0-fold after 12 h (Fig. S15a). Next, we examined our RNA-seq datasets to determine if genes in the PSR network were induced. Indeed, 72 of the 179 genes in the PSR were upregulated in roots following exposure to 20 µM juglone (Fig. 5b). While the expression of most genes was induced within the first 12 h of exposure and attenuated by 24 h, several genes remained upregulated by 5 d. Within the PSR subset, we additionally observed induction of numerous 26S proteasome regulatory particle subunits, including most lid (RPN5/6/7/8/9/10/11/12) and base (RPN1/2, RPT1-6) components, and the accessory factor PA200 (Fig. S15b). Thus, these data indicate that juglone exposure triggers proteasome biogenesis and the induction of protein quality control mechanisms associated with the PSR.

**Fig. 5.**
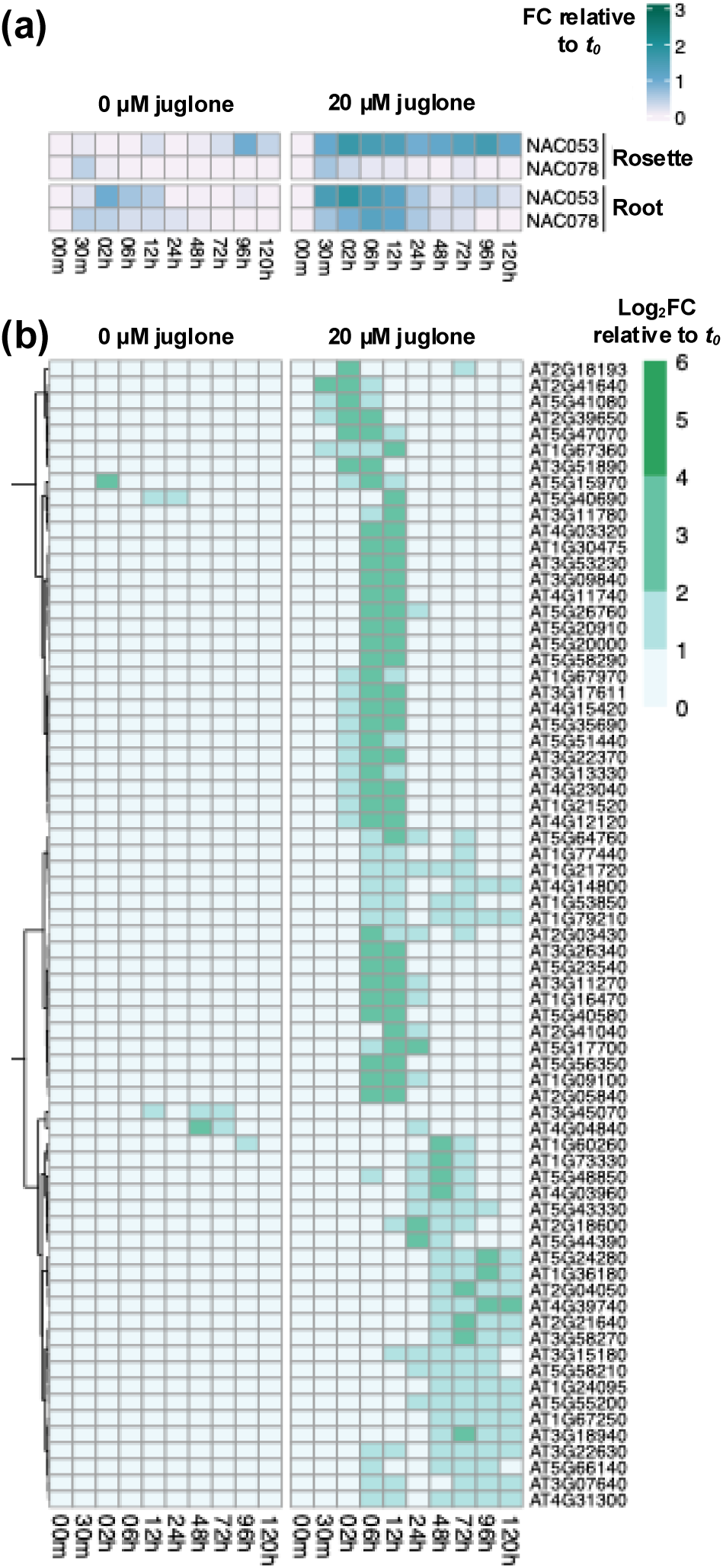
The proteasome stress regulon (PSR) is induced by juglone. (a) Representation of juglone-induced changes in transcript levels of *NAC53* (At3g10500) and *NAC78* (At5g04410) over the RNA-seq time series. Color represents the fold-change (FC) in gene expression level relative to *t_0_*. Significant (FDR<0.01) differential expression of *NAC53* in rosettes and *NAC78* in the roots was observed. (b) Heatmap representation of data from the RNA-seq experiment showing the expression of 72 out of 179 genes comprising the proteasome stress regulon (Gladman et al., 2016) that were upregulated (log₂FC ≥ 1) in Arabidopsis roots at one or more time points between 0.5 and 120 hours after exposure to 20 µM juglone, relative to *t₀*. Expression patterns following transfer to mock control media (0 µM juglone) are shown for comparison. Note that FC was used to depict expression in (a) while log_2_FC was used in (b) to better illustrate changes in expression.

### Proteomics analysis reveals broad proteome remodeling in response to juglone

To better understand how juglone proteotoxicity alters the proteome, we conducted untargeted DIA-based proteomics on whole 12-d-old Arabidopsis plants 12 h after exposure to 20 µM juglone or mock treatment. Approximately 6200 proteins were identified at a 1% false discovery rate under each treatment. Principal component analysis (PCA) revealed distinct proteomic shifts at both time points (Fig. S16a). In juglone-treated plants, numerous proteins were found to be more abundant, including several GST subunits, heat shock proteins (HSPs), and stress-responsive proteins, such as members of the MOS4-associated splicing complex, which is involved in response to heat stress (Endo *et al*., 2023) and the dehydrins (Szlachtowska & Rurek, 2023) (Fig. 6a). KEGG pathway enrichment analysis revealed “glutathione metabolism” and “protein processing in the ER” among the enriched categories in proteins that accumulated under juglone stress (Fig. S16b). Conversely, certain oxidative repair cycle proteins, including thioredoxin (Trx) and monodehydroascorbate reductase (MDAR), were less abundant (Fig. 6a). Considering Trx relies on active-site cysteines, this may indicate that Trx is alkylated by juglone or the Trx pool shifts toward a highly oxidized state, triggering its degradation. Although MDAR does not use GSH directly, it is part of the ascorbate-glutathione cycle, which depends on both NADPH and GSH to maintain reduced ascorbate. The reduced abundance of MDAR may reflect the depletion of GSH and disruption of the ascorbate-glutathione cycle.

**Fig. 6.**
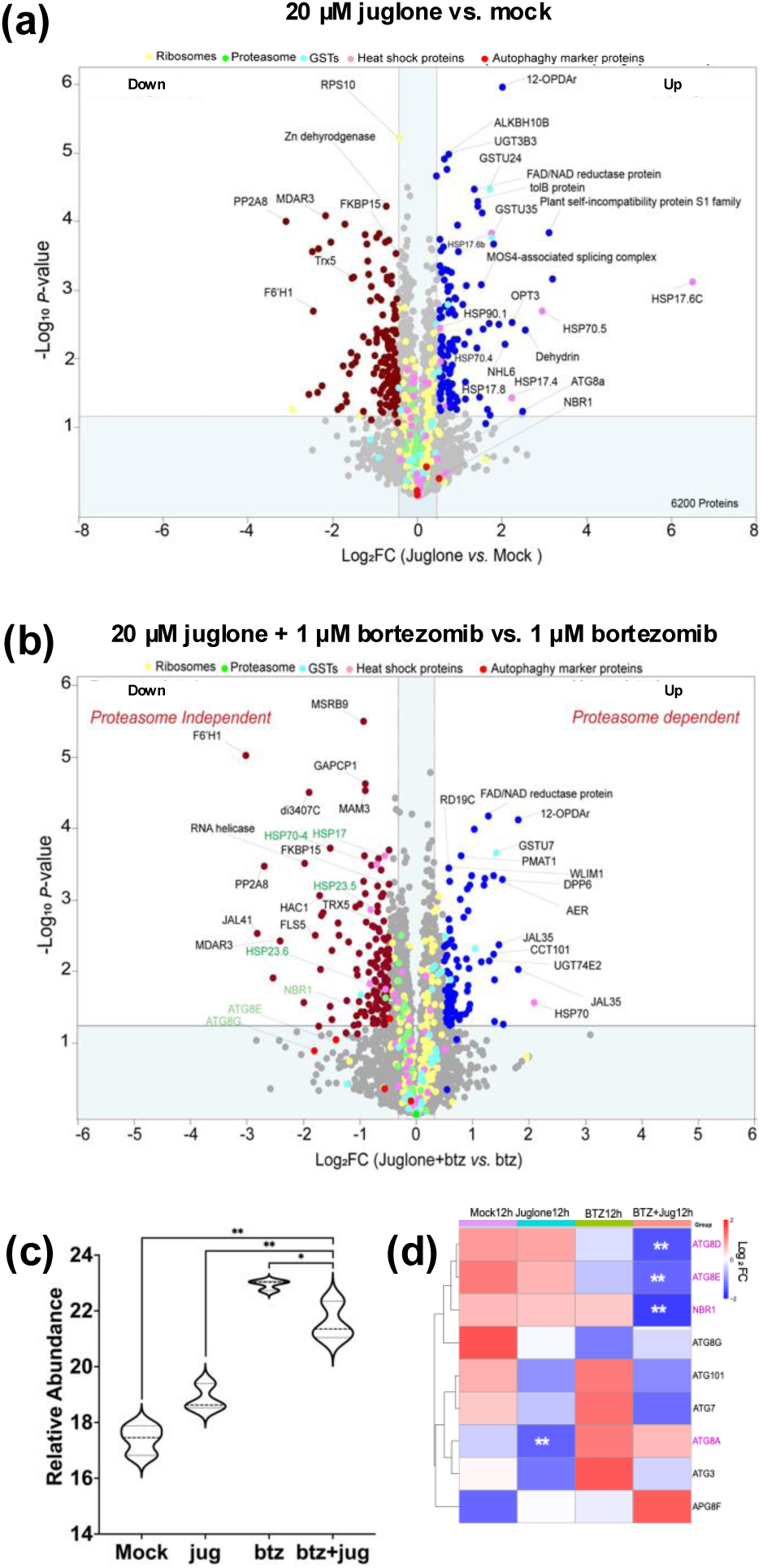
Proteomic responses to juglone from experiments with whole 12-d-old Arabidopsis plants 12 h after treatment. (a) Volcano plot of differentially accumulated proteins following a 12-h exposure to 20 µM juglone compared to mock treatment. Approximately 6,200 proteins are displayed. Accumulated proteins (“up”), passing both *p*-value and fold-change cutoffs as indicated on the plot, are in blue; proteins with statistically significant lower abundances (“down”) are in maroon. Selected key proteins and families are labeled and color-coded as indicated. (b) Volcano plot of differentially accumulated proteins following a 12-h exposure to 20 µM juglone and 1 µM of the proteasome inhibitor bortezomib (btz) compared to btz alone. Approximately 6,240 proteins are displayed. Accumulating proteins (“up”), passing both *p*-value and fold-change cutoffs as indicated on the plot, are in blue; proteins with statistically lower abundances (“down”) are in maroon. Selected key proteins and families are labeled and color-coded as indicated. (c) Violin plots show the protein intensity levels of NBR1. NBR1-MS1 levels from data-dependent acquisition are shown. The NBR1 level increases under both juglone and bortezomib treatment. The decrease in the level of NBR1 upon combined treatment of juglone and bortezomib, indicates a shuttling of substrates for degradation via autophagy. (d) Heat map showing changes in autophagy marker proteins across the root and seedling proteomic datasets under juglone treatment.

In response to juglone, proteasome subunits, *NAC53* and *NAC78*, and many PSR genes were upregulated at the transcriptional level (Fig. 5 and Fig. S15). To further investigate the role of the proteasome, we used bortezomib, a reversible proteasome inhibitor that binds to core protease subunits, leading to inactivation of the proteasome and the accumulation of ubiquitinated proteins (Tan *et al*., 2018). We treated 12-d-old Arabidopsis plants with 20 µM juglone and 1 µM bortezomib for 12 hours and compared their proteome to plants treated with 1 µM bortezomib alone. Principal component analysis revealed distinct proteomic profiles among plants treated with juglone and bortezomib, bortezomib alone, juglone alone, or mock (Fig. S17). Several proteasome subunits were significantly upregulated following bortezomib treatment compared to mock controls, confirming its effectiveness in Arabidopsis. We next examined the combinatorial effects of juglone and bortezomib. Proteomic analysis revealed a distinct expression pattern, with volcano plots showing downregulation of several core autophagy components (Fig. 6b). In line with this trend, HSPs, which were upregulated by juglone treatment alone (Fig. 6a), appeared downregulated under the combined treatment (Fig. 6b). Notably, HSPs are recognized autophagy substrates during plant stress responses. We then examined whether the ubiquitin-binding autophagy receptor NEIGHBOR OF BRCA1 gene 1 (NBR1) was also affected. In plants, NBR1 has a role in selective autophagy where it promotes the accretion of ubiquitylated proteins into larger condensates that are encapsulated by autophagosomes for macroautophagic clearance in some, but not all, autophagy events (Jung *et al*., 2020; Zhang & Chen, 2020; Lee *et al*., 2023). Using both DIA and DDA datasets (MS1-level quantifications), we found a notable decrease in the level of NBR1 in the combined juglone and bortezomib treatment (Fig. 6c). Note, lower NBR1 levels act as a proxy for higher autophagic flux since NBR1 itself is an autophagy substrate and is degraded when autophagy is active. This suggests that juglone-induced proteotoxic substrates are redirected to autophagic degradation when the availability of functional proteasome is limited. To investigate this further, we examined known autophagy substrates, core components, and receptors (Fig. 6d). Previous work has demonstrated that in Arabidopsis, proteasome abundance is controlled by an autophagy-related gene (ATG)-8 mediated mechanism (Marshall *et al*., 2015). Indeed, the abundances of several ATG8 proteins from different clades decreased, including ATG8D, ATG8E, and ATG8A, indicating active shuttling of these substrates for degradation via autophagy-mediated vacuolar degradation. Overall, the transcriptomics and proteomics data presented are consistent with juglone inducing proteotoxic stress in Arabidopsis with clearance of damaged proteins through the PSR, engaging both the proteasome and autophagy.

## DISCUSSION

Here, we report a systems-level understanding of how Arabidopsis responds to the phytotoxic allelochemical juglone. Integration of transcriptomics and proteomics with biochemical experiments revealed that juglone widely modifies thiol groups in proteins and GSH, leading to the coordinated activation of responses in redox buffering, in the metabolism of glutathione, sulfur, and cysteine, and in protein quality control (i.e., upregulation of chaperones, the proteasome, and autophagy components). These responses reflect juglone’s general chemical nature as a quinone to redox cycle and alkylate with nucleophiles (O’Brien, 1991; Bolton & Dunlap, 2017). The ability of juglone to induce reactive oxygen and nitrogen species is well documented, e.g., (Babula *et al*., 2014; Chen *et al*., 2015; Kurtyka *et al*., 2016), and its propensity to covalently modify thiols was first observed decades ago (Sturdik *et al*., 1983). Nonetheless, juglone’s mechanism of action is still considered as an inhibitor of specific targets, like the plasma membrane H⁺-ATPase in plants (Jose & Gillespie, 1998; Dayan & Duke, 2014) and Pin1-type peptidyl-prolyl isomerases in bacteria and various eukaryotes (Hennig *et al*., 1998; Chao *et al*., 2001; Yao *et al*., 2001). At the time of writing this manuscript, juglone is commercially available at Sigma-Aldrich (St. Louis, MO, USA; catalog number 420120) under the description of being “a highly selective cell-permeable, irreversible inhibitor” of peptidyl-prolyl isomerases, and is used on this basis as a cell culture inhibitor in many studies. There is a need to correctly frame observed biological responses to juglone in the context of its chemical properties. Our study supports a model aligned with juglone’s dual chemistry and indiscriminate interactions with multiple targets, via its propensity to redox cycle, its electrophilicity, or both, leading to the depletion of GSH, protein misfolding, and the a priori widespread loss of cellular enzyme activities. Whether it be a plant or animal, juglone is likely to elicit cytotoxicity through a combination of ROS, thiol depletion, and as a general disrupter of proteostasis. While we specifically examined cysteine thiols, the electrophilicity of juglone likely extends to attack by other cellular nucleophiles, including the amino groups on L-lysine, the N-termini of proteins, and the nucleobase nitrogen and oxygen atoms in DNA and RNA.

In plants, pathways to synthesize 1,4-naphthoquinones have convergently evolved many times (Meyer *et al*., 2021). Our study raises new questions about how the chemistries of different 1,4-naphthoquinones contribute to their relative abilities to alkylate proteins in vivo, disrupt redox systems, and elicit multi-target stress responses. It also highlights gaps in understanding about how the different chemical properties of naphthoquinones translate to their ecological roles in allelopathy, deterring herbivores, and as antimicrobial compounds. These questions draw comparisons to the chemistries and functions of other plant electrophiles, such as isothiocyanates released from the hydrolysis of glucosinolates. Like the 1,4-naphthoquinones, isothiocyanates are electrophilic and capable of modifying cellular nucleophiles, including cysteine thiols (Sun *et al*., 2025). Sulforaphane depletes GSH, induces cell death, and primes defenses during the hypersensitive response in Arabidopsis (Andersson *et al*., 2015). The electrophilicity of isothiocyanates also underlies the biochemical mechanism responsible for chemical defenses against certain insect herbivores (Schramm *et al*., 2012; Jeschke *et al*., 2016). It therefore seems that electrophilic small molecules could represent a broadly evolved strategy for mediating plant-biotic interactions.

The PSR has been well studied in the context of induced proteasome limitation (Gladman *et al*., 2016), but its importance in maintaining protein quality control in response to environmental stresses is only just beginning to be understood (Langin *et al*., 2023; Yu *et al*., 2024). Our study reveals that soon after juglone exposure, plants activate the PSR, including the upregulation of dozens of downstream targets of NAC53 and NAC78 involved in protein folding, proteasome assembly, and autophagy (Gladman *et al*., 2016) (Fig. 5). This suggests the direct involvement of both the 26S proteasome and selective autophagy pathways for clearing damaged substrates. Moreover, the accumulation of HSPs and the downregulation of NBR1 and ATG proteins during combined juglone stress and proteasome inhibition (Fig. 6) resemble proteotoxicity responses observed during heat stress (Thirumalaikumar *et al*., 2021). Heat is among the few other environmental cues known to simultaneously activate both the proteasome and autophagy pathways in plants (Ohama *et al*., 2017; Raffeiner *et al*., 2023). This suggests that juglone also leads to protein misfolding and aggregation on a significant level. Given that HSPs themselves can become substrates of autophagy when overwhelmed (Thirumalaikumar *et al*., 2021), our similar observation when combining juglone treatment with the proteasome inhibitor bortezomib (Fig. 6b) implies that autophagy may act as a backup clearance mechanism when proteasome capacity is limited, especially during continual exposure to high levels of juglone.

It is important to note that in our experiments, juglone concentrations were approximately 10-100 times lower than what plants experience near black walnut trees (De Scisciolo *et al*., 1990). Thus, continual and higher exposures to juglone in nature may elicit a proteotoxic load that far exceeds the capacity of the basal proteostasis machinery and thus prove a broad and efficient deterrent to seedling establishment in proximity to the black walnut trees. Arrested root growth and foraging in response to juglone (Fig. 1b) likely reflect a constant and futile diversion of resources from growth and development toward repairing and turning over damaged proteins and other small molecules and macromolecules. Several NAC transcriptions, known for their roles as central regulators of developmental processes and stress acclimation (Xiong *et al*., 2025), showed various degrees of upregulation in roots in response to juglone (Fig. S18). These provide a starting point for examining the transcriptional networks linking redox and proteotoxic cues from juglone to the reprogramming of root growth. Of note is *NAC103*, which plays a role in relaying ER stress signals from bZIP60 to downstream genes involved in the unfolded protein response (UPR) (Sun *et al*., 2013). The induction of *NAC103* is especially high at later time points (48-120 h), while that of *NAC78* attenuates (Fig. S18). This may indicate a transition from an early PSR-dominated response to a longer-term strategy in which ER stress and UPR pathways help to maintain the integrity of the root proteome under persistent juglone exposure.

Another outstanding question is how juglone and other allelopathic naphthoquinones affect quinone signaling networks in plants. Plant roots are continuously bathed in quinones, whether they are encountered as allelochemicals intentionally released from other organisms or as chemical byproducts of oxidized phenolics or other cellular debris in the rhizosphere. It remains to be investigated if or how juglone and other naphthoquinones interact with the membrane-bound receptor CANNOT RESPOND TO DMBQ 1 (CARD1) (Laohavisit *et al*., 2020), also termed HYDROGEN-PEROXIDE-INDUCED CA^2+^ INCREASES 1 (HPCA1) (Wu *et al*., 2020). The CARD1/HPCA1 receptor has an extracellular domain that protrudes into the apoplast, containing four cysteine residues that, when oxidized by quinones or hydrogen peroxide, trigger an intracellularly linked kinase domain to initiate a calcium signaling cascade leading to stress responses (Fichman *et al*., 2022; Zhang *et al*., 2025). Examination of our RNA-seq data indicates that juglone does not induce expression of *CARD1/HPCA1* or its homologs (Fig. S19). However, induction of *CARD1/HPCA1* is not necessarily expected even if juglone can trigger the receptor, given its ubiquitous expression throughout plant development (Wu *et al*., 2016) and role in sensing via post-translational regulation by cysteine oxidation. At the same time, however, several *CARD1*-dependent marker genes are upregulated by juglone (Fig. S19). Moreover, inhibition of root growth by juglone (Fig. 1b and Fig. S1) (Meyer *et al*., 2021) resembles the CARD1-dependent inhibition by 2,6-dimethoxy-1,4-benzoquinone (DMBQ) and hydrogen peroxide (H_2_O_2_) (Zhang *et al*., 2025). Given that Arabidopsis root growth is also sensitive to the absolute amount of glutathione rather than to changes in the glutathione redox potential (Safi *et al*., 2026), juglone-driven glutathione depletion provides a plausible mechanistic link between quinone perception, activation of CARD1/HPCA1-dependent stress pathways, and the arrest of root meristem activity.

Our study underscores the need to investigate how black walnut and other plants that produce 1,4-naphthoquinones prevent autotoxicity. While a lack of molecular specificity makes 1,4-naphthoquinones a broadly effective ecological strategy, their promiscuity presents a biochemical challenge that requires the simultaneous evolution of robust mechanisms to mitigate self-inflicted damage. Given that the majority of juglone in *Juglans* species is glycosylated (Duroux *et al*., 1998; McCoy *et al*., 2018), conjugation and storage could be serving a role in preventing autotoxicity. However, this strategy requires preferential glycosylation on the naphthoquinone ring (e.g., at the C4 position), like that in the structure of hydrojuglone glucoside (1,5-dihydroxy-4-naphthalenyl-β-D-glucopyranoside). Glycosylation at the C5 position of hydrojuglone or juglone (Fig. 1a) would result in the naphthoquinone ring remaining available for redox cycling and alkylation. The glycosylation mechanism would also rely on spontaneous hydrolysis of the deployed conjugate in the rhizosphere, or enzymatic hydrolysis by free enzymes in the soil or upon uptake by other organisms. In addition, walnut has apoplastic glucosidases capable of hydrolyzing hydrojuglone glucoside to juglone (Duroux *et al*., 1998), which could conceivably serve as a control point for activating the quinone outside of the cell. Such a compartmentalized activation strategy parallels the mechanism of the glucosinolate-myrosinase system, in which a conjugated electrophile is stored in an inert form and enzymatically activated later.

This study also has implications and raises concerns for some of the translational applications proposed for juglone (Islam & Widhalm, 2020). For example, juglone has long been considered a promising natural product-based herbicide candidate. However, despite its strong phytotoxicity (Fig. 1b, Fig. S1), juglone lacks several traits desirable in a commercial herbicide. It lacks molecular specificity, it acts at high concentrations, and it requires prolonged exposure. Its promiscuity also raises concerns about off-target effects on crops, soil microbes, and beneficial fungi. Similarly, the promiscuity of naphthoquinones should raise questions about the regular consumption of herbal supplements containing high levels of bioactive naphthoquinones. While the naphthoquinone scaffold may provide promising chemistry for agricultural and medicinal applications, without improved specificity, targeted delivery, or controlled activation, these compounds have the potential to disrupt proteostasis and redox balance in non-target organisms, including humans.

## ACKNOWLEDGEMENTS

This work was supported by United States Department of Agriculture-National Institute of Food and Agriculture (USDA-NIFA) Agriculture and Food Research Initiative project numbers IND00065150G to K.V. and J.R.W. and IND00165252G to G.W.M., and USDA-NIFA Hatch Project numbers 177845 to J.R.W. and 1013620 to K.V. The authors thank the Purdue Metabolite Profiling Facility and Purdue Proteomics Facility associated with the Bindley Bioscience Center for the metabolomics and proteomics analyses. The authors thank the National Institutes of Health for the timsTOF HT awarded to Purdue (1S10OD032364-01A1).

## Figure Legends

**Supporting Information Fig. S1.**
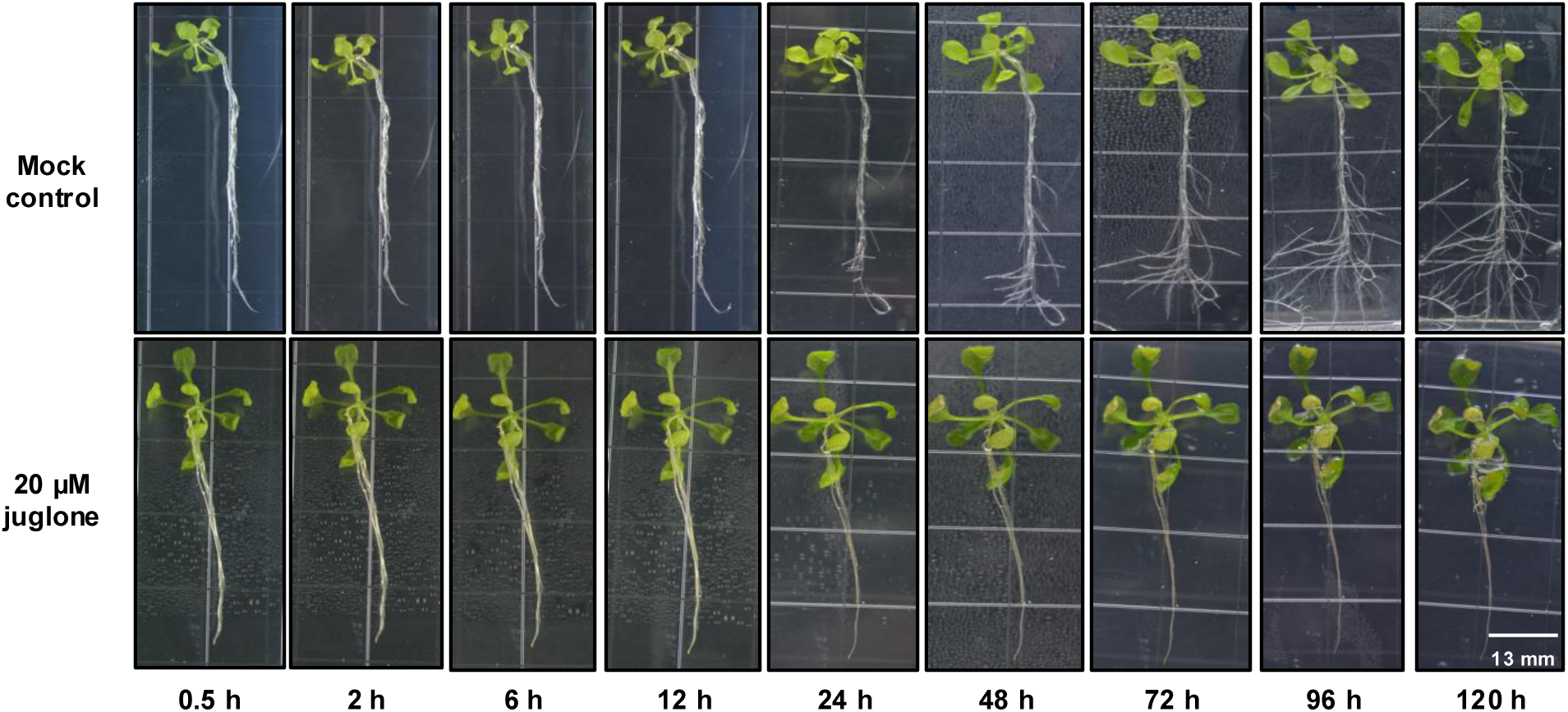
Sampling time course for RNA-seq analysis. Images show abaxial view of representative 12-d-old Arabidopsis plants grown on ½ MS and transferred to new media containing 0 µM (mock) or 20 µM juglone and imaged from 0.5 h to 5 d (120 h). At each time point indicated, roots and rosettes were harvested separately for RNA-seq (n=3 biological replicates with organs from ≥4 plants per replicate). By 24 h after transfer, mock-treated roots started to initiate lateral root formation with continued growth and foraging over the course of the experiment, while juglone-treated roots did not. Grid squares are 13x13 mm.

**Supporting Information Fig. S2.**
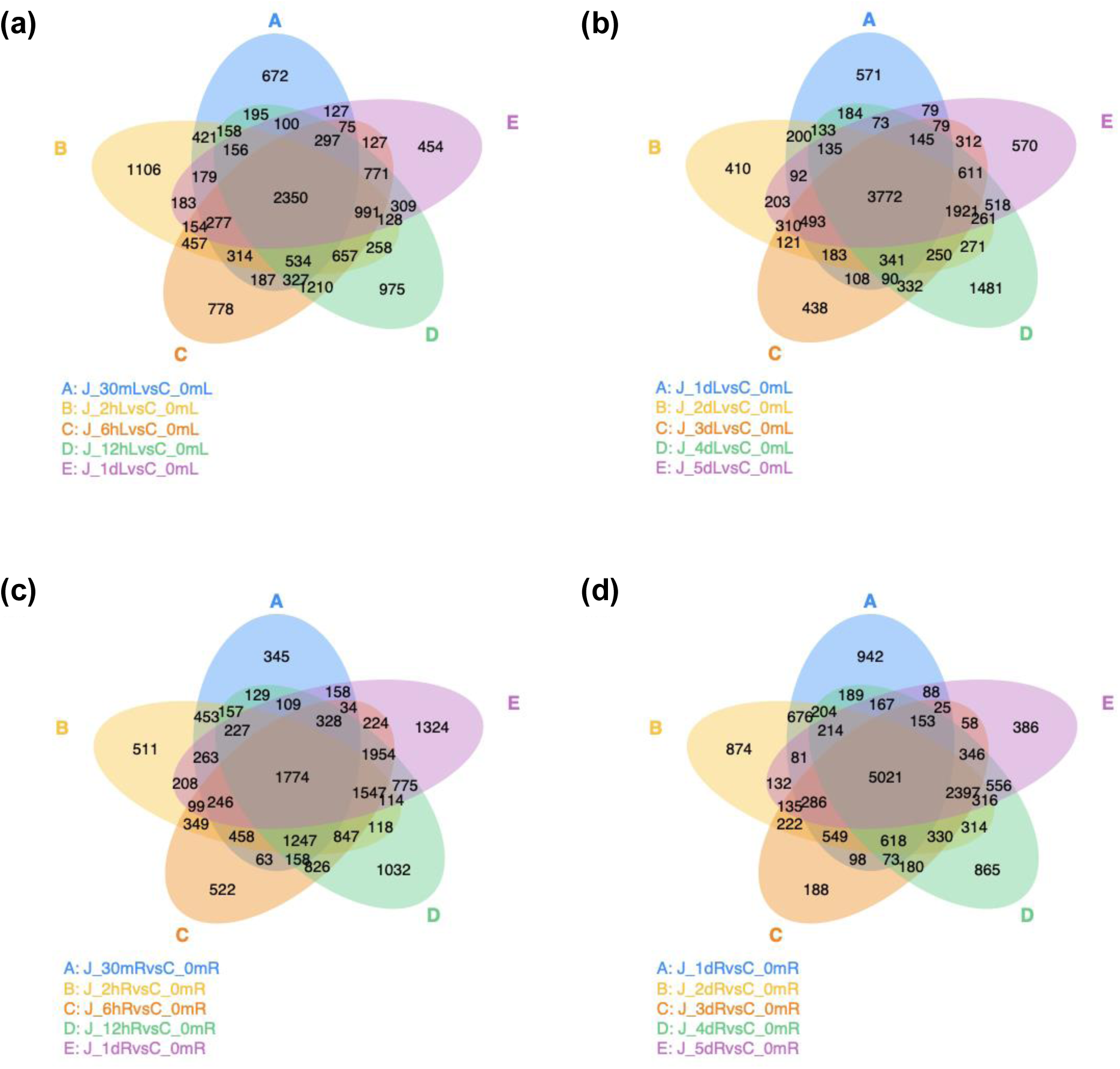
Venn diagrams showing the overlap of differentially expressed genes across time points in juglone-treated Arabidopsis rosettes and roots compared to organs from untreated plants sampled at *t_0_* (before transferring plants to treatment media). Differentially expressed genes (DEGs) identified in rosettes (a,b) and roots (c,d) across early (a,c) and late (b,d) time points. Early response includes 0.5 h, 2 h, 6 h, 12 h, and 24 h, while late response includes 24 h, 48 h, 72 h, 96 h, and 120 h.

**Supporting Information Fig. S3.**
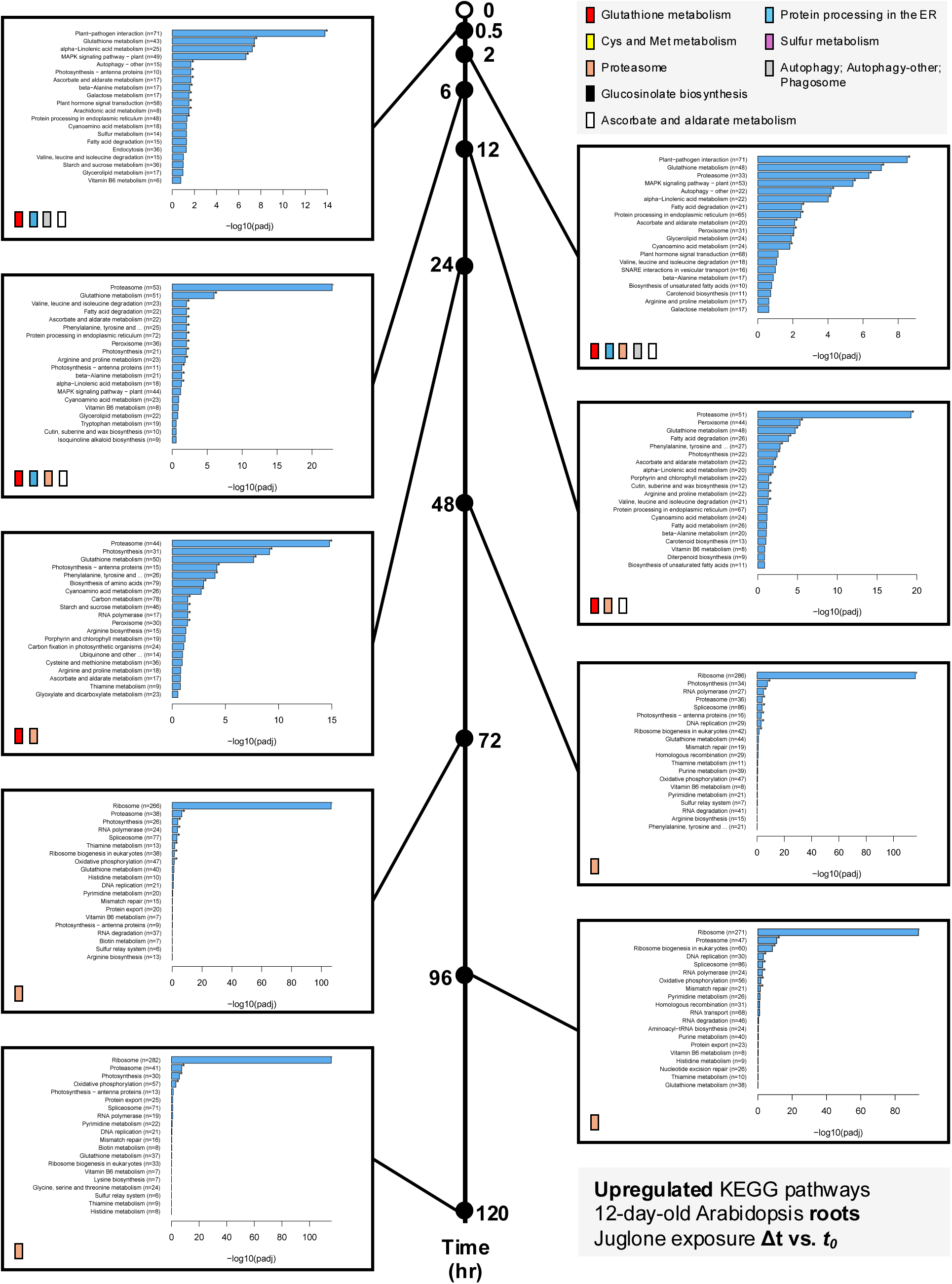
Significantly upregulated KEGG pathways in 20 µM juglone-exposed roots of 12-day-old Arabidopsis plants compared to untreated roots sampled at *t_0_* (before transferring plants to treatment media). Pathways discussed in the text are highlighted by the legend in the top right corner.

**Supporting Information Fig. S4.**
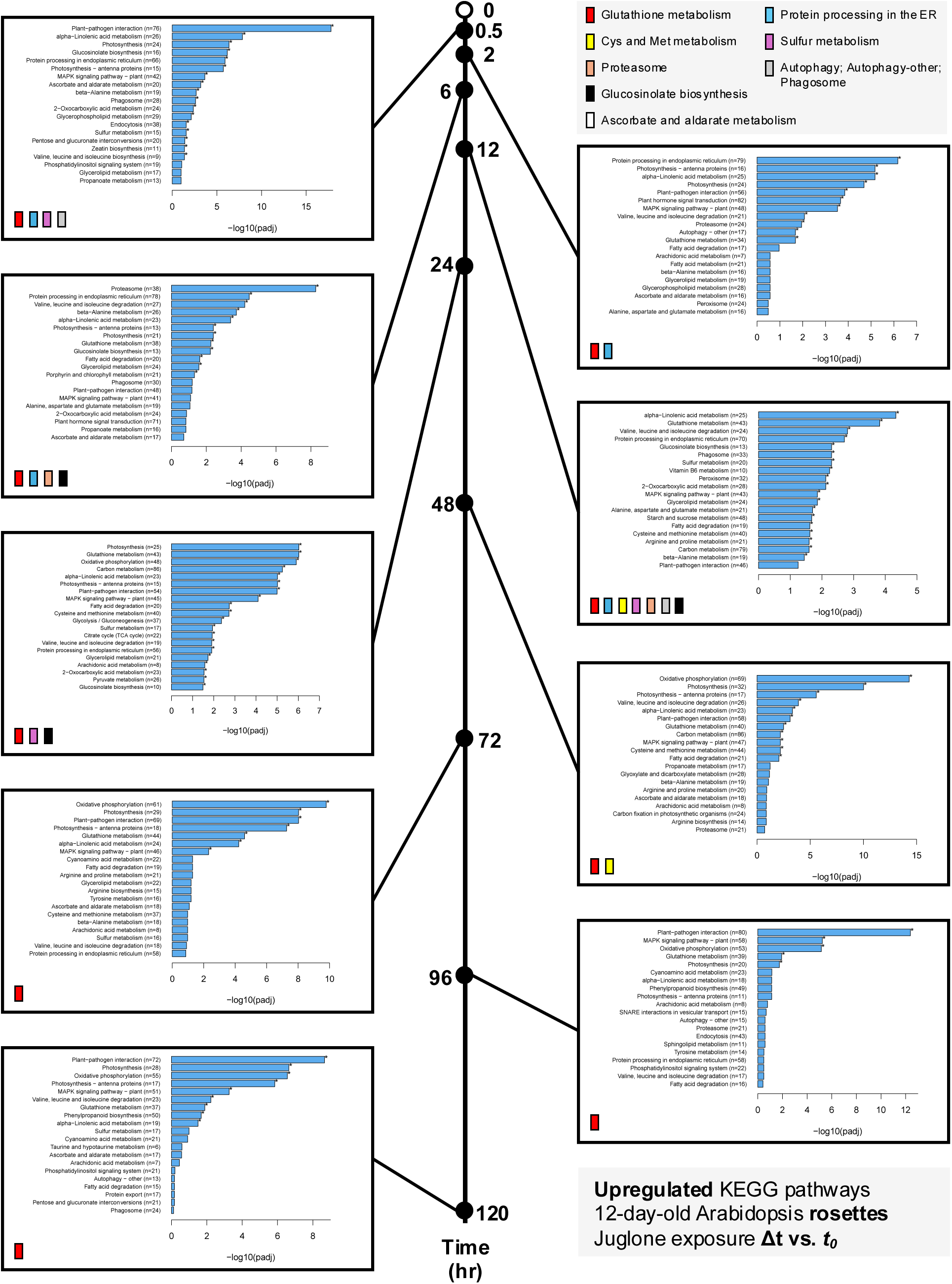
Significantly upregulated KEGG pathways in 20 µM juglone-exposed rosettes of 12-day-old Arabidopsis plants compared to untreated rosettes sampled at *t_0_* (before transferring plants to treatment media). Pathways discussed in the text are highlighted by the legend in the top right corner.

**Supporting Information Fig. S5.**
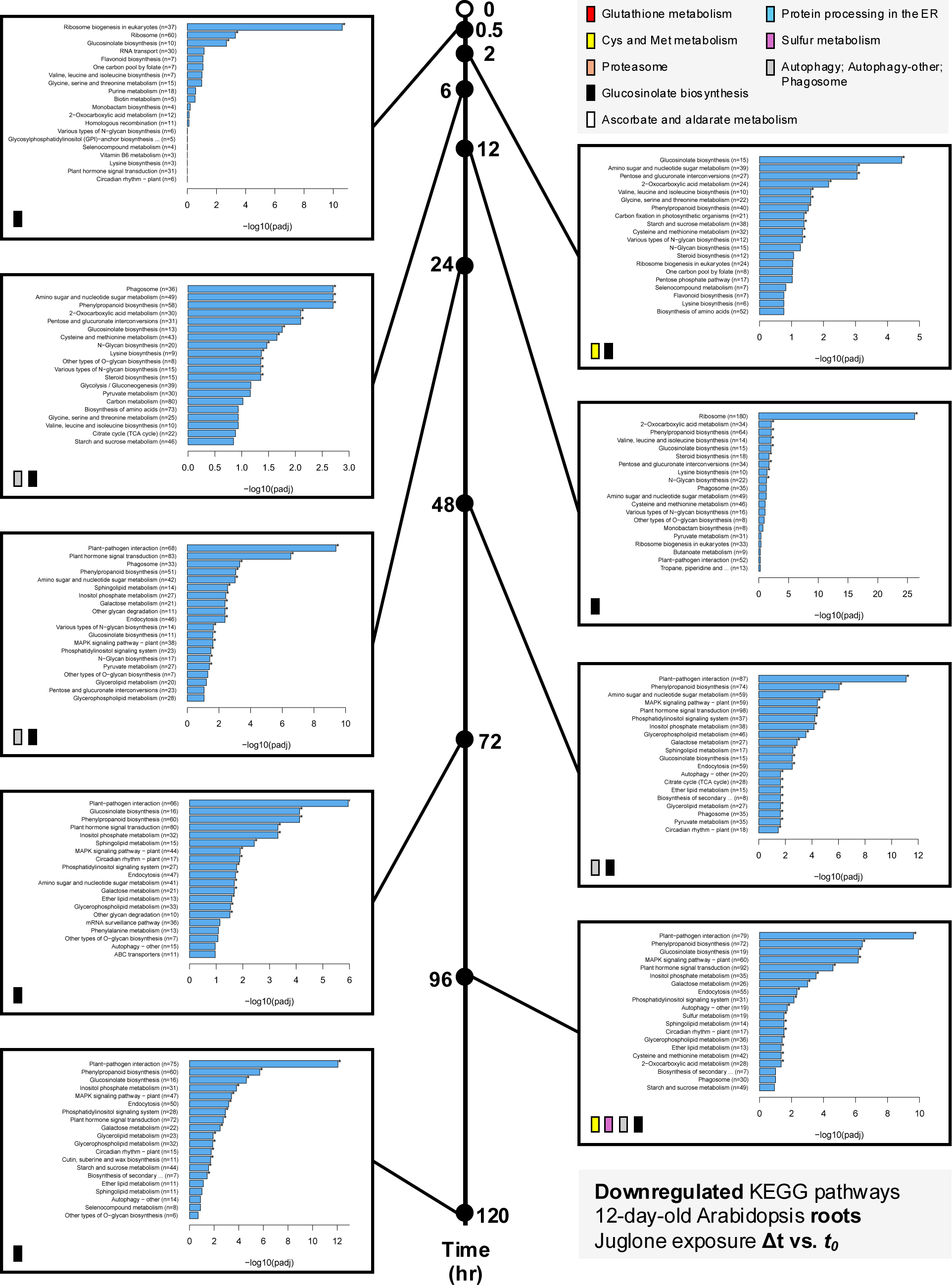
Significantly downregulated KEGG pathways in 20 µM juglone-exposed roots of 12-day-old Arabidopsis plants compared to untreated roots sampled at *t_0_* (before transferring plants to treatment media). Pathways discussed in the text are highlighted by the legend in the top right corner.

**Supporting Information Fig. S6.**
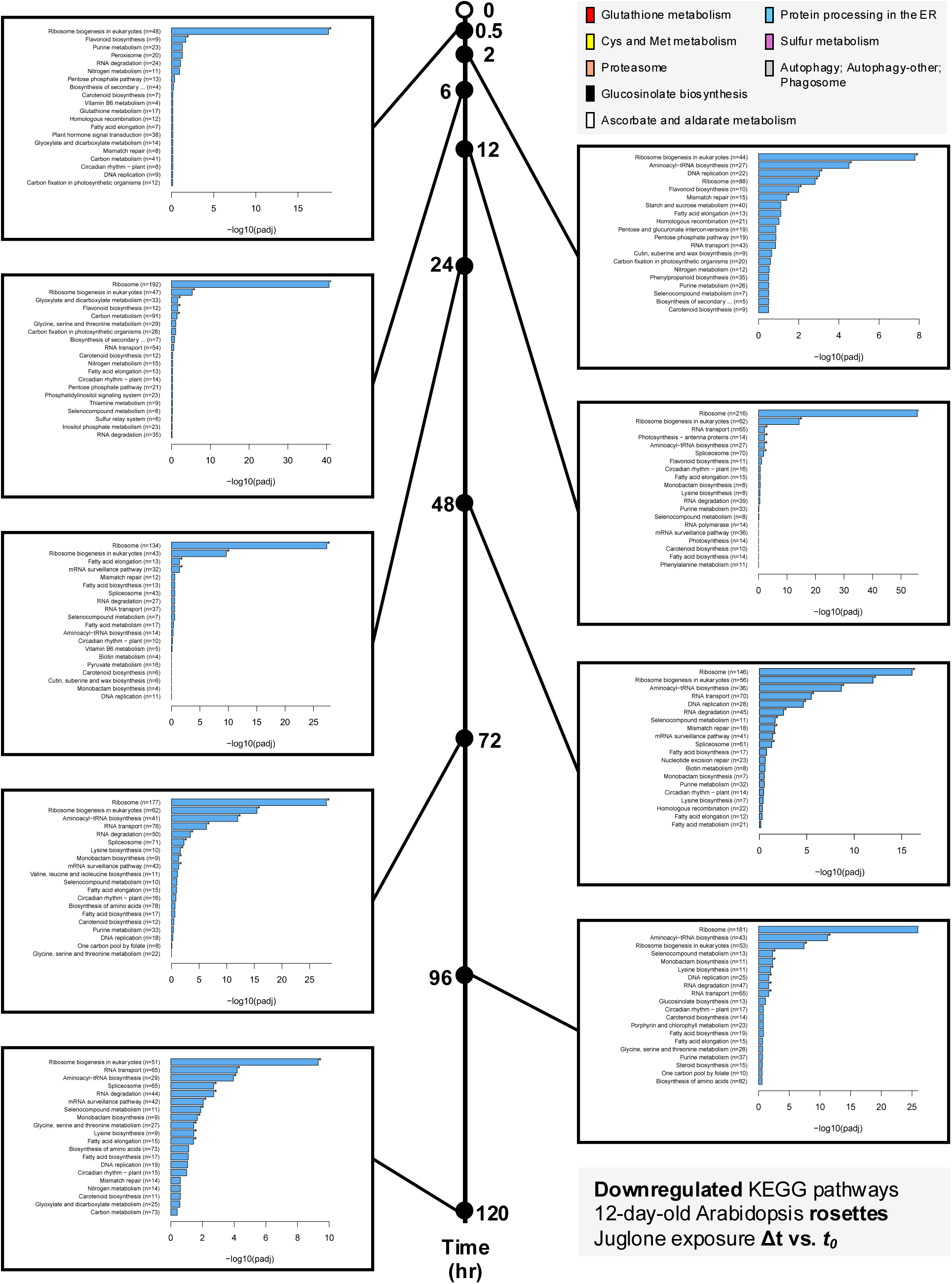
Significantly downregulated KEGG pathways in 20 µM juglone-exposed rosettes of 12-day-old Arabidopsis plants compared to untreated rosettes sampled at *t_0_* (before transferring plants to treatment media). Pathways discussed in the text are highlighted by the legend in the top right corner.

**Supporting Information Fig. S7.**
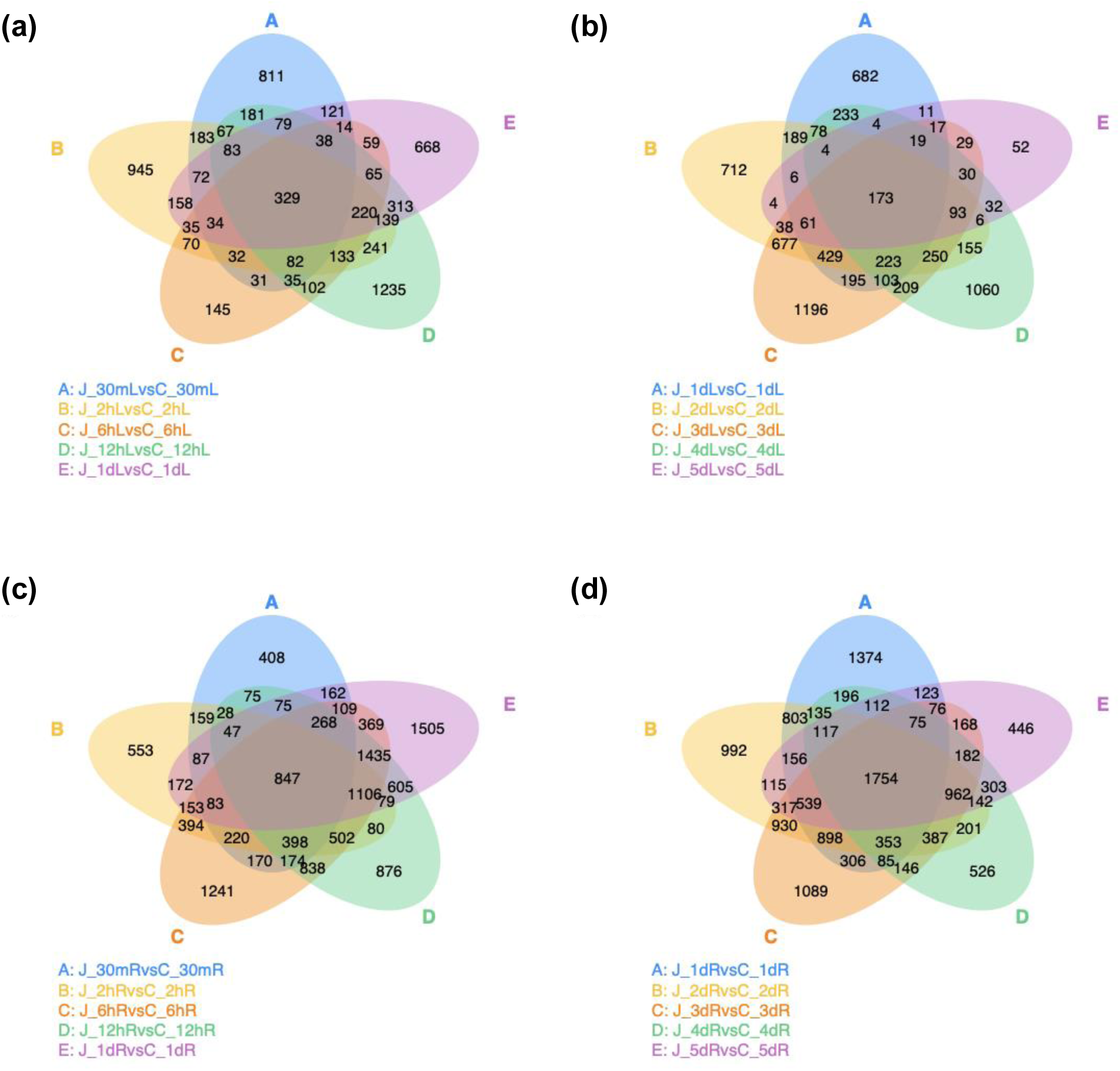
Venn diagrams showing the overlap of differentially expressed genes across time points in juglone-treated Arabidopsis rosettes and roots. Differentially expressed genes (DEGs) identified in rosettes (a,b) and roots (c,d) across early (a,c) and late (b,d) time points. Comparisons are between organs samples from juglone-treated and mock-treated plants at each time point. Early response includes 0.5 h, 2 h, 6 h, 12 h, and 24 h, while late response includes 24 h, 48 h, 72 h, 96 h, and 120 h.

**Supporting Information Fig. S8.**
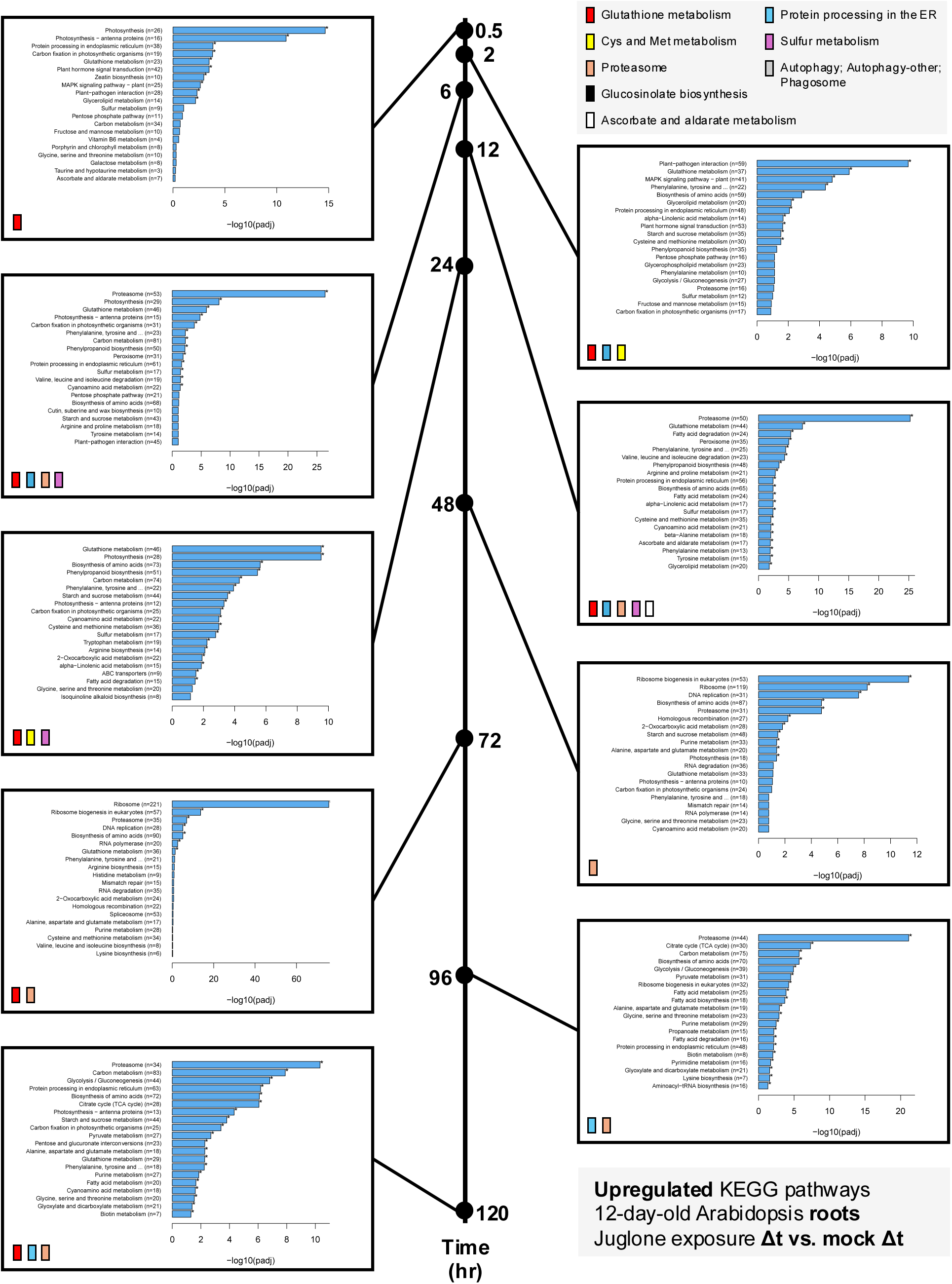
Significantly upregulated KEGG pathways in 20 µM juglone-exposed roots of 12-day-old Arabidopsis plants compared to mock-treated roots sampled at the same time (Δt) after transferring plants to treatment media. Pathways discussed in the text are highlighted by the legend in the top right corner.

**Supporting Information Fig. S9.**
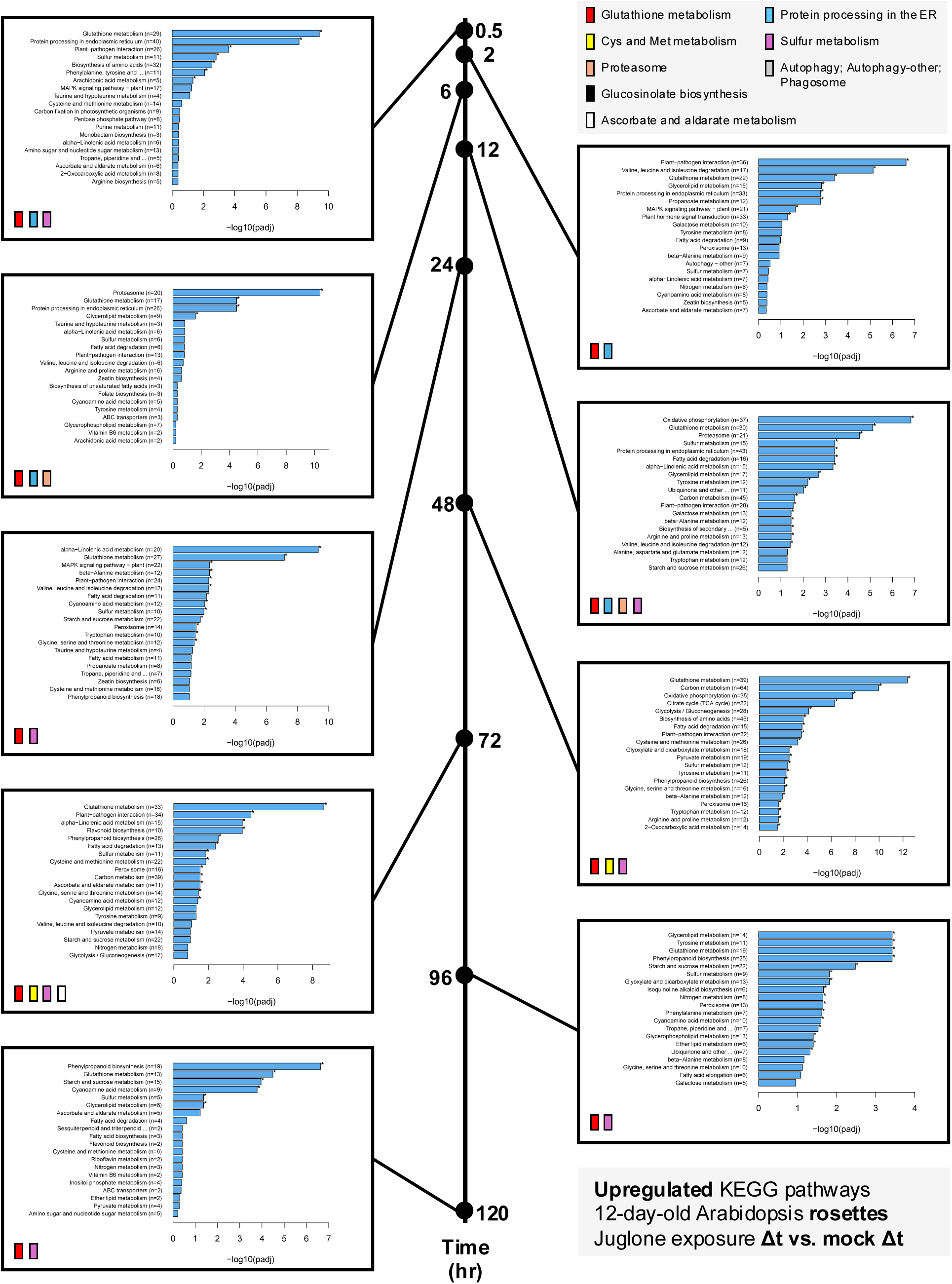
Significantly upregulated KEGG pathways in 20 µM juglone-exposed rosettes of 12-day-old Arabidopsis plants compared to mock-treated rosettes sampled at the same time (Δt) after transferring plants to treatment media. Pathways discussed in the text are highlighted by the legend in the top right corner.

**Supporting Information Fig. S10.**
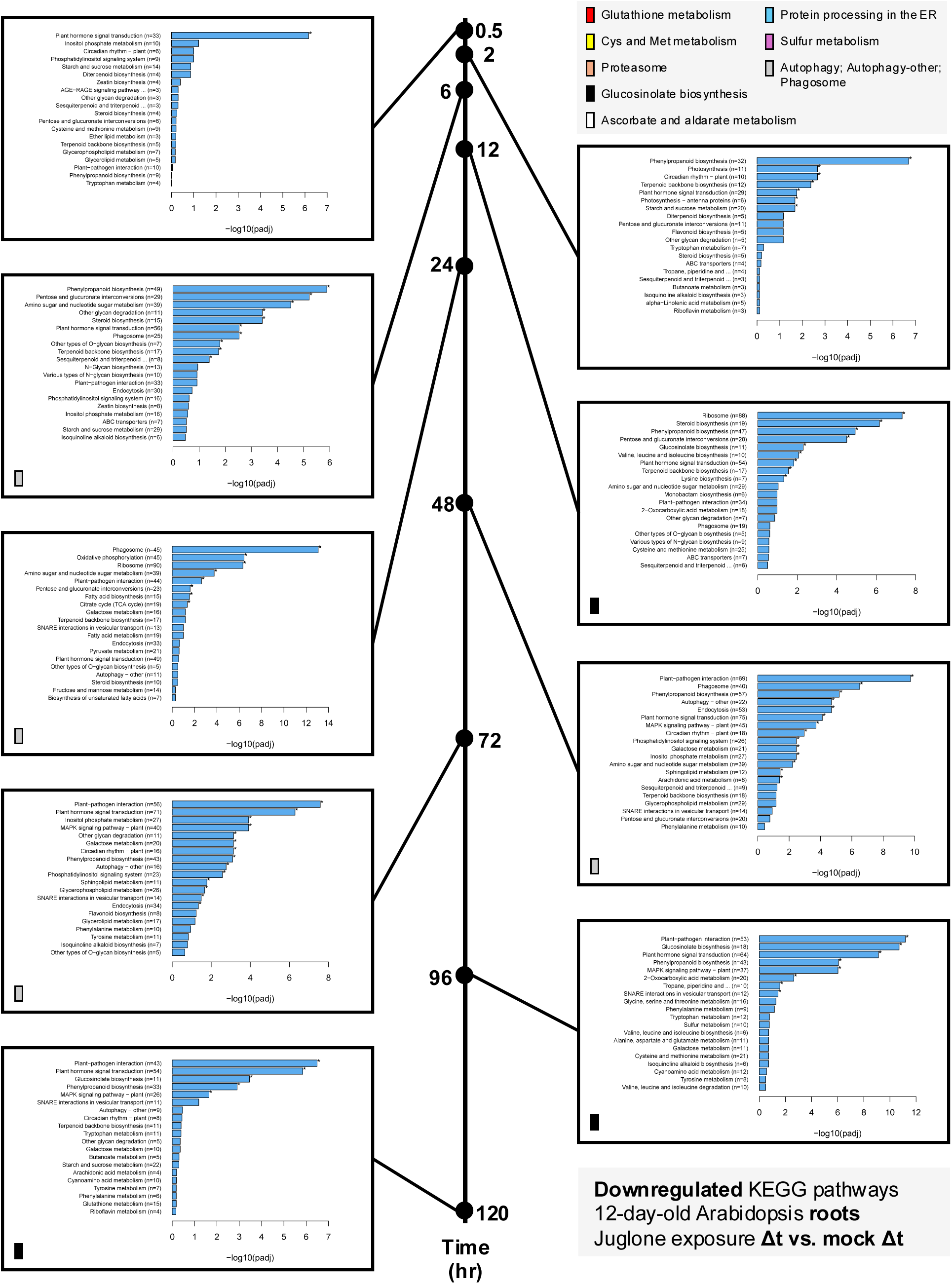
Significantly downregulated KEGG pathways in 20 µM juglone-exposed roots of 12-day-old Arabidopsis plants compared to mock-treated roots sampled at the same time (Δt) after transferring plants to treatment media. Pathways discussed in the text are highlighted by the legend in the top right corner.

**Supporting Information Fig. S11.**
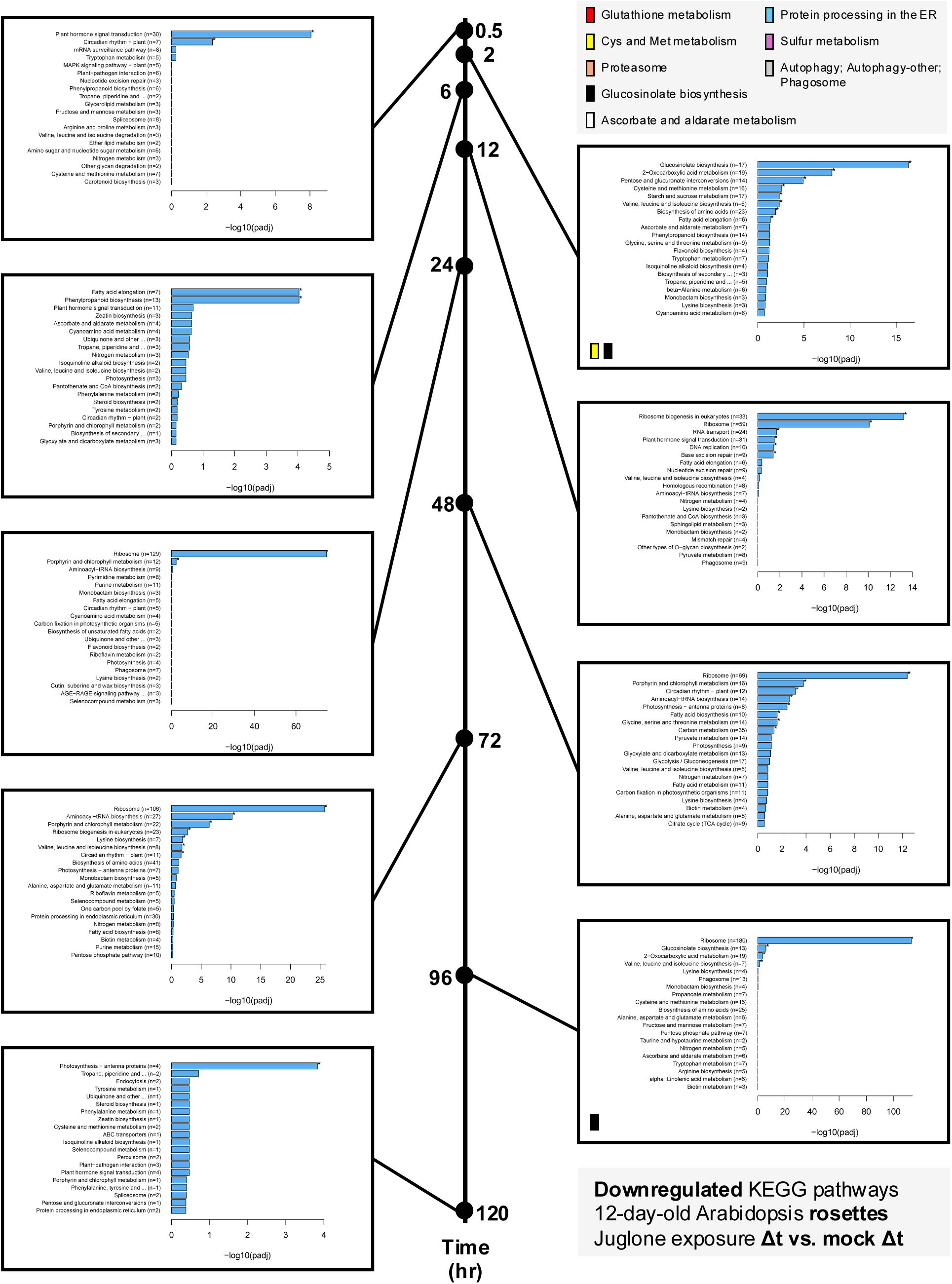
Significantly downregulated KEGG pathways in 20 µM juglone-exposed rosettes of 12-day-old Arabidopsis plants compared to mock-treated rosettes sampled at the same time (Δt) after transferring plants to treatment media. Pathways discussed in the text are highlighted by the legend in the top right corner.

**Supporting Information Fig. Fig. S12.**
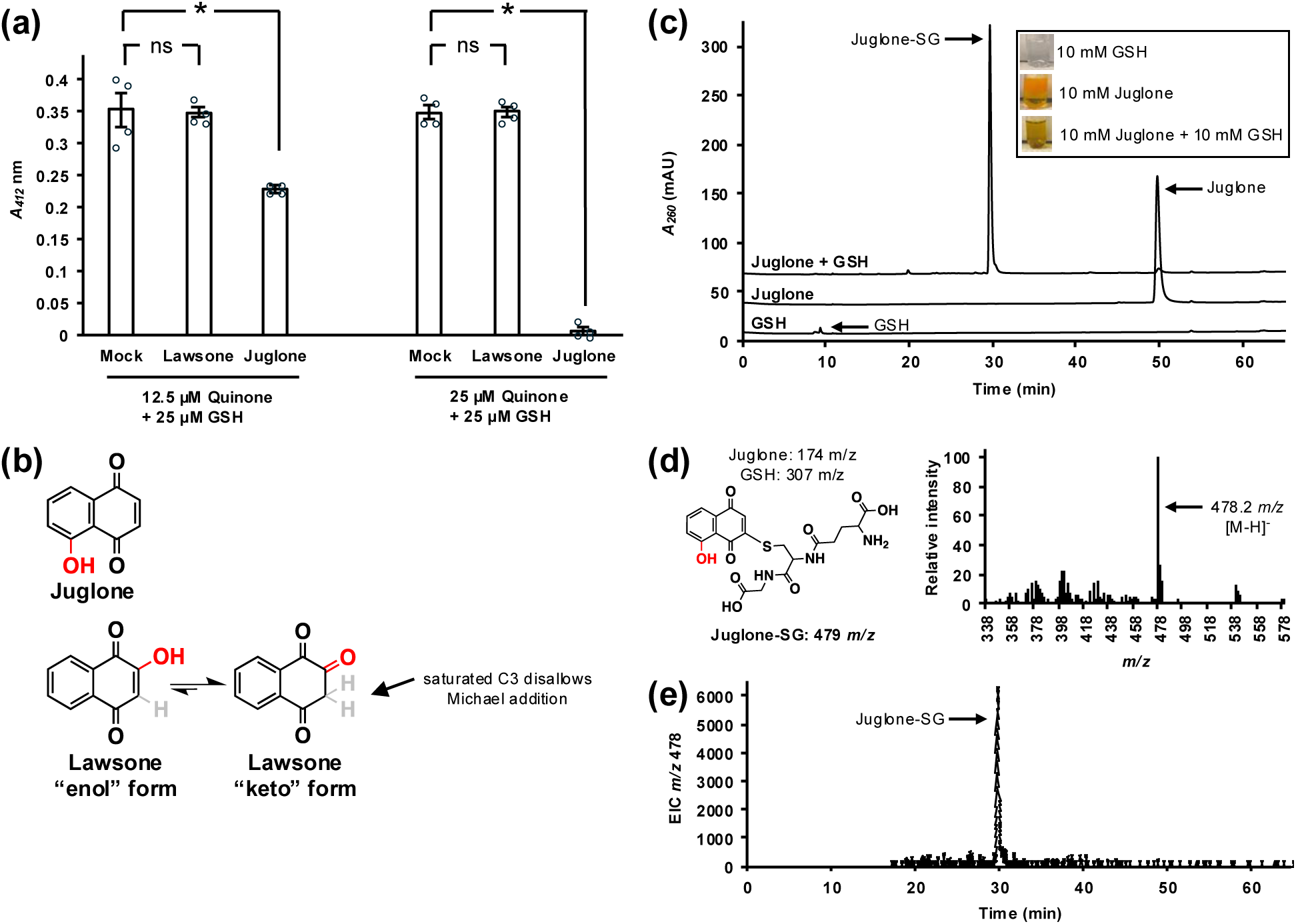
Juglone spontaneously forms adducts with reduced glutathione. (a) Quantification of free thiols after incubation of 25 µM lawsone, juglone, or mock with 12.5 or 25 µM reduced glutathione (GSH). Bars show mean ± SE (n=4); *p < 0.05, Student’s *t*-test; ns, not significant. (b) Chemical structures of juglone and of lawsone in its enol and keto forms. Tautomerization favors the keto form, which reduces the electrophilicity of lawsone at C3. (c) HPLC-DAD analysis with detection at *A*_260_ nm showing elution profiles of juglone, GSH, and a reaction mixture of the two. A new peak eluting at 29.8 min appears only in the mixture and was identified as the juglone-GSH conjugate, juglone-SG. Inset photos show visible color differences of the reaction mixtures prior to HPLC analyses. (d) LC-MS analysis of the corresponding peak at 29.78 min revealed a major ion, *m/z* 478.2, which is consistent with the expected mass of juglone-SG formed via spontaneous Michael addition of GSH to juglone. (e) The extracted ion chromatogram (EIC) for *m/z* 478 confirms that the conjugate elutes at at 29.8 min.

**Supporting Information Fig. S13.**
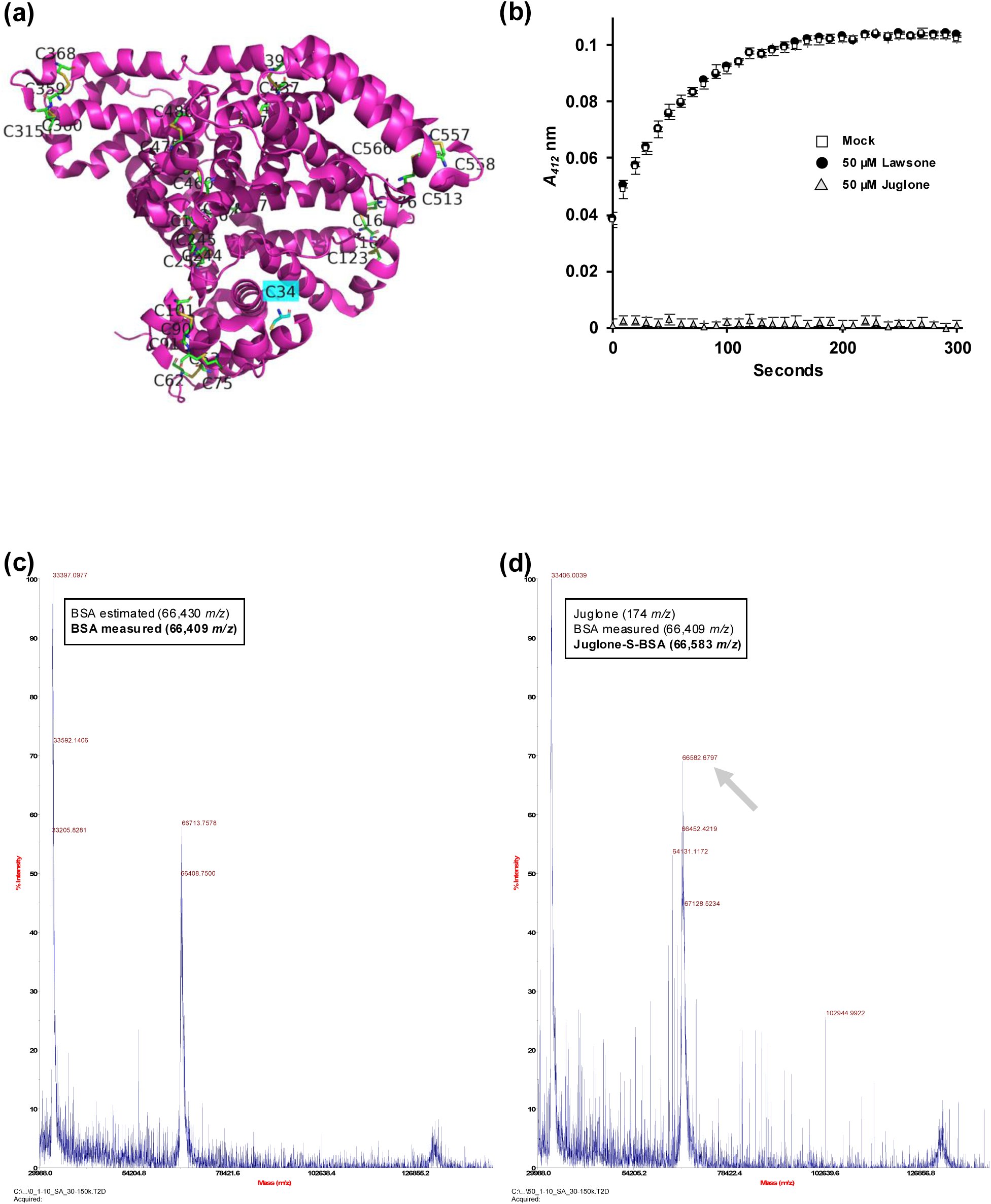
Juglone covalently modifies accessible cysteine residues in bovine serum albumin (BSA). (a) Homomeric structure of BSA (Protein Data Bank ID: 4F5S) with its lone accessible free cysteine, C34, highlighted in cyan. (b) Quantification of free thiols in BSA after incubation with 50 µM juglone, lawsone, or mock for 5 min. Data show mean ± SE (n = 3). (c,d) MALDI-TOF mass spectrometry of BSA alone (c) or incubated with juglone (d). A new peak (arrow) corresponding to a mass increase of ∼174 Da in the juglone-treated sample is consistent with the covalent addition of one juglone molecule (174 *m/z*) to BSA (66,583 *m/z* vs. unmodified 66,409 *m/z*).

**Supporting Information Fig. S14.**
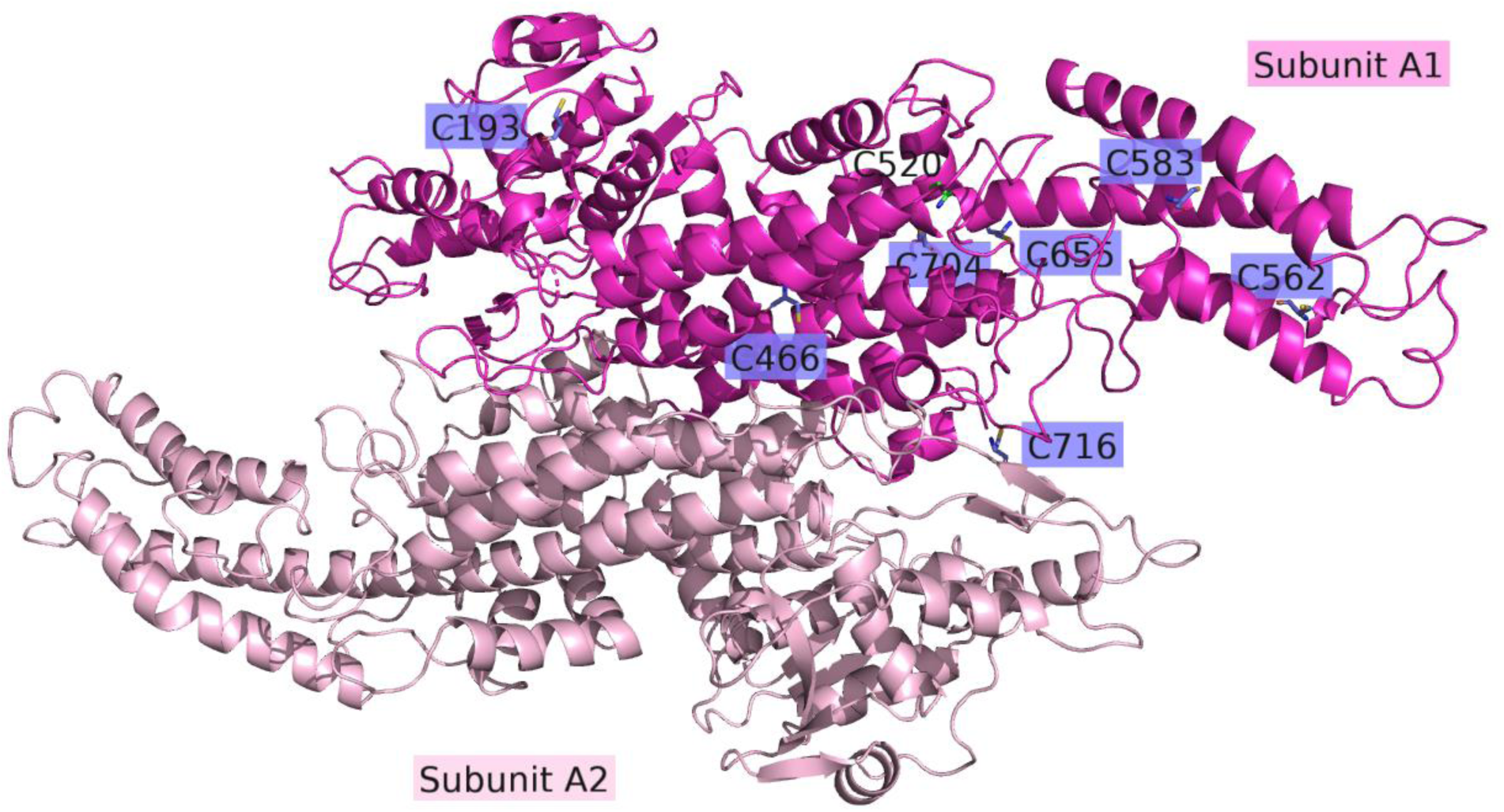
The structure of *Petroselinum crispum* (parsley) phenylalanine ammonia-lyase (PAL) homodimer (Protein Data Bank ID: 1W27). The parsley PAL homomer has eight cysteine residues, seven of which are likely to be located on the surface of the protein (indicated in blue) based on the predicted structure.

**Supporting Information Fig. S15.**
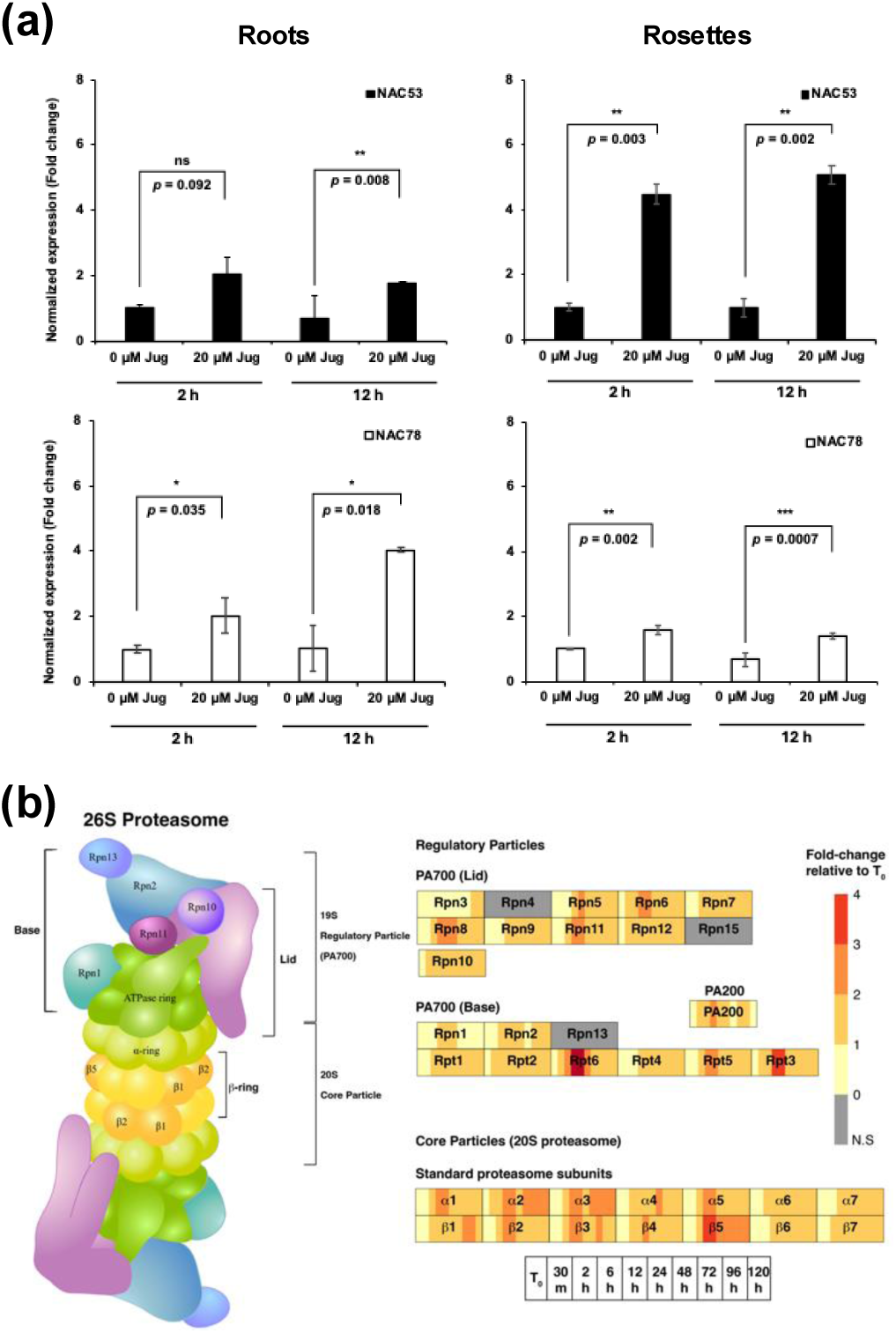
The proteasome stress regulon (PSR) is induced by juglone. (a) RT-qPCR validation of *NAC53* and *NAC78* expression in roots and rosettes of 12-d-old Arabidopsis 2 and 12 h after exposure to 20 µM juglone or mock treatment. Expression values were determined from total RNA and calculated relative to *ACT2*. Values are mean ± SE (n=3 with 3 plants per replicate); *p < 0.05, Student’s *t*-test versus respective mock control. (b) Schematic representation of the 26S proteasome showing regulatory particles (PA700 lid and base) and the 20S core particle, with adjacent heatmaps summarizing juglone-induced transcript changes over time for the corresponding subunit genes in roots (log₂fold-change relative to t₀). Colors indicate mean expression fold-change across biological replicates from the RNA-seq time course. Data in (b) and are from the RNA-seq experiment described in Supplementary Fig. S1.

**Supporting Information Fig. S16.**
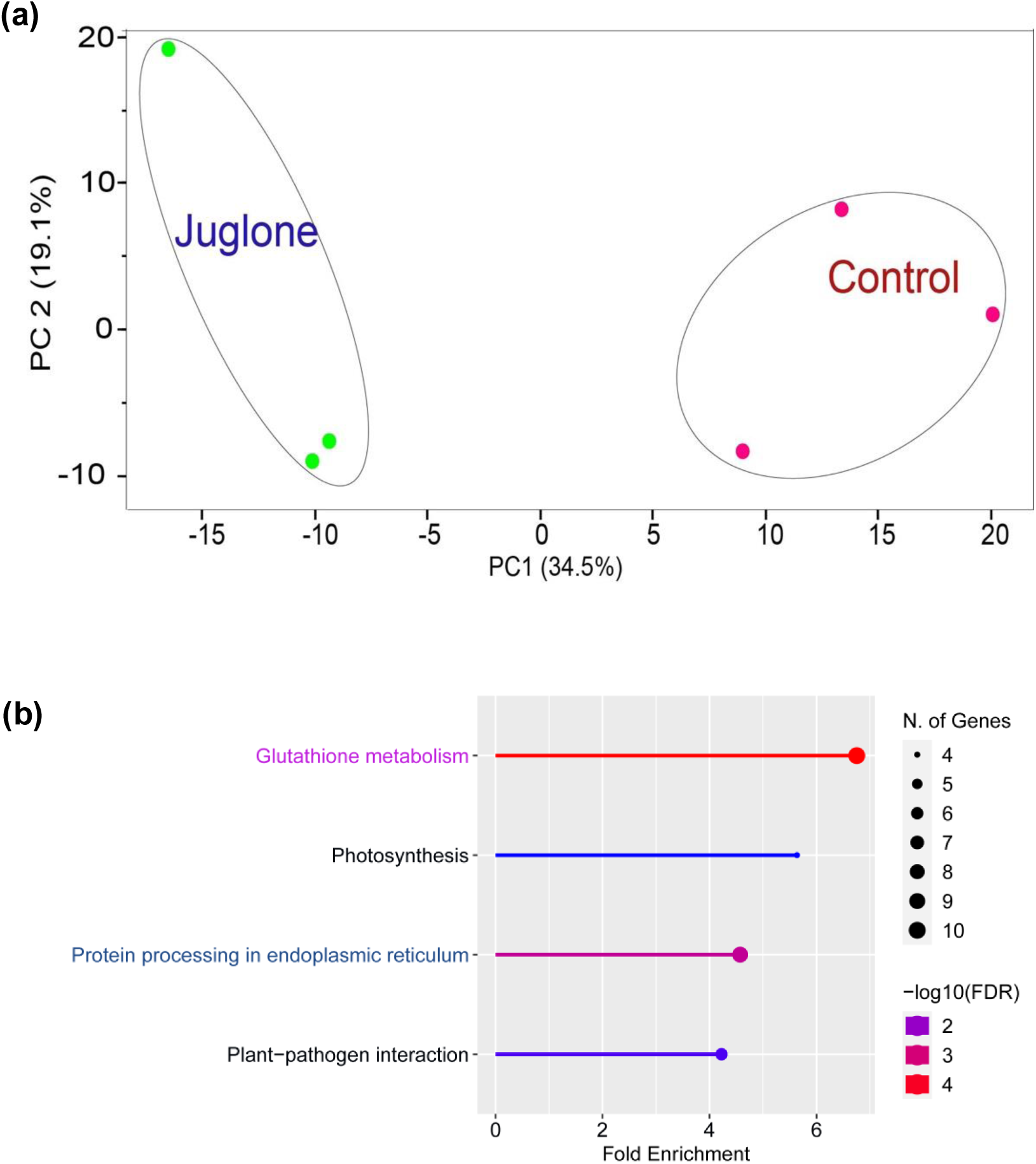
Juglone treatment alters the Arabidopsis proteome. (a) Principal component analysis (PCA) of the seedling proteomic samples. Log-transformed protein intensities from data-independent acquisition (DIA) were plotted for proteomes sequences from whole 12-d-old Arabidopsis plants 12 h following treatment with 0 µM (control) or 20 µM juglone. PC1 and PC2 values are indicated. (b) KEGG pathway enrichment analysis of pathways over-represented in juglone-exposed Arabidopsis. The enrichment was calculated using ShinyGO v82 using background genes.

**Supporting Information Fig. S17.**
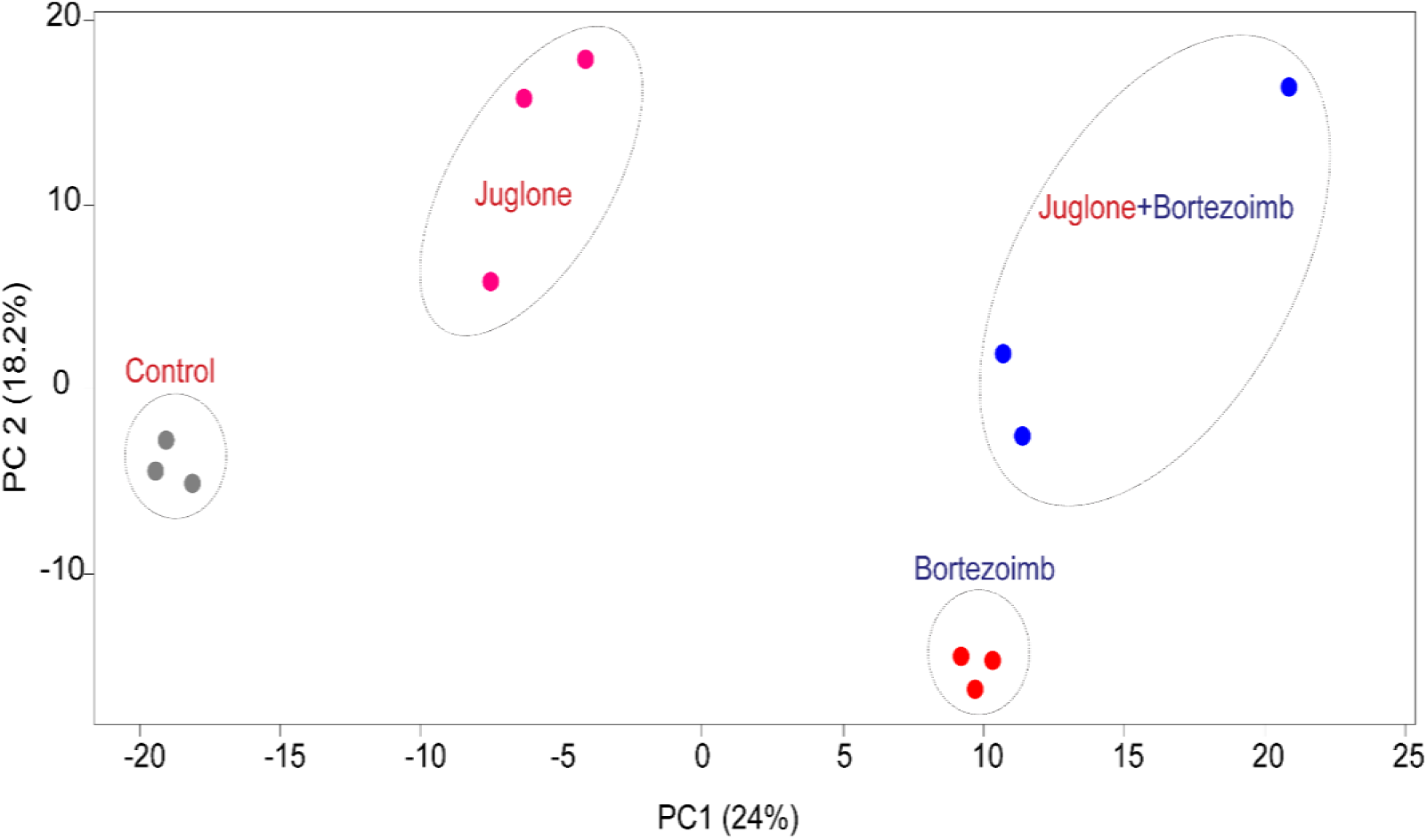
Principal component analysis (PCA) of Arabidopsis proteomic samples following a 12-h exposure to mock treatment, 20 µM juglone, 20 µM juglone + 1 µM of the proteasome inhibitor bortezomib (btz), or btz alone. Log-transformed protein intensities from data-independent acquisition (DIA) were plotted. PC1 and PC2 values are indicated.

**Supporting Information Fig. S18.**
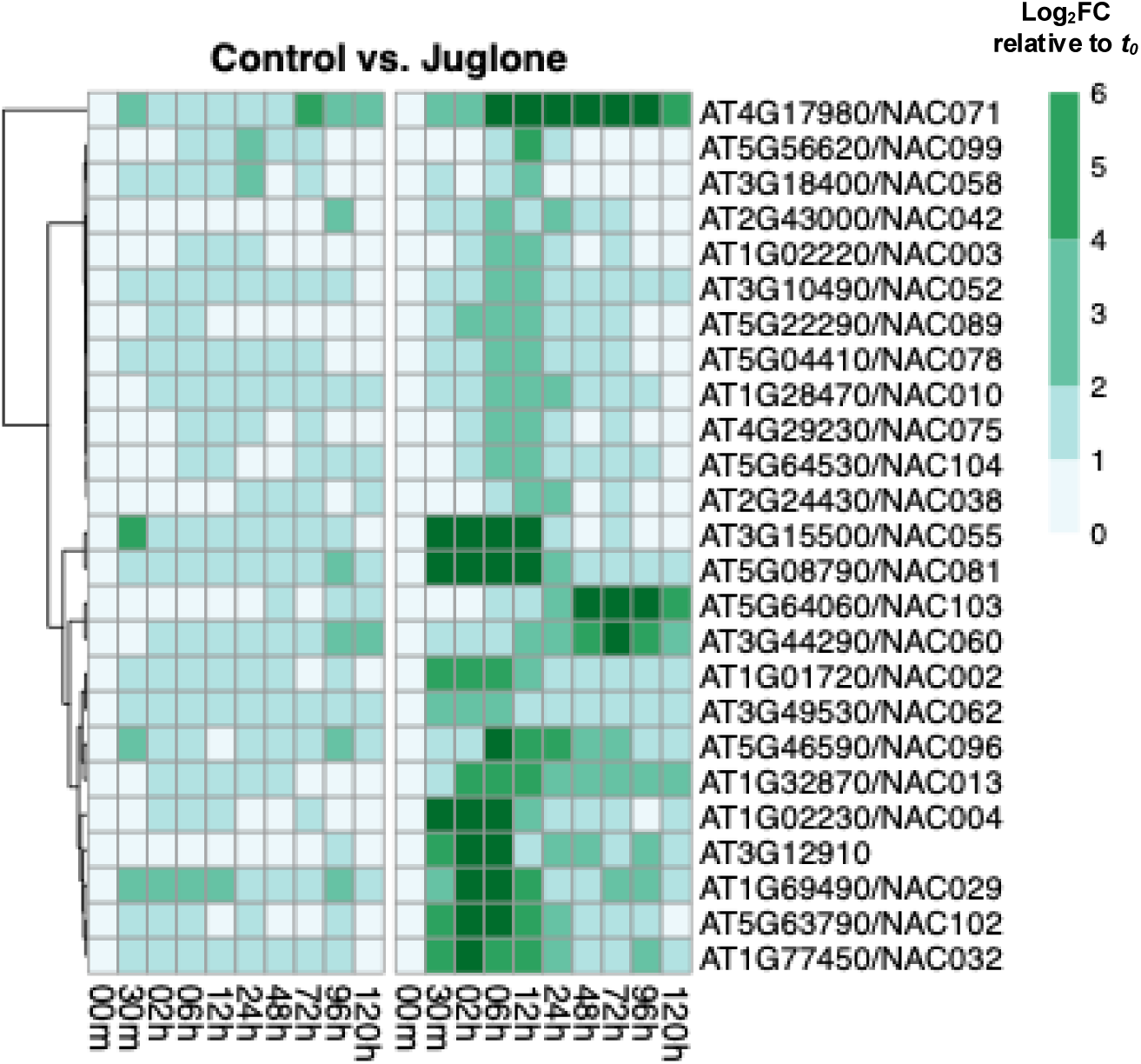
Heatmap representation of data from the RNA-seq experiment showing the expression of 25 NAC transcription factor genes that were upregulated (log₂FC ≥ 1) in Arabidopsis roots at one or more time points between 0.5 and 120 hours after exposure to 20 µM juglone, relative to *t_0_*. Expression patterns following transfer to mock control media (0 µM juglone) are shown for comparison.

**Supporting Information Fig. S19.**
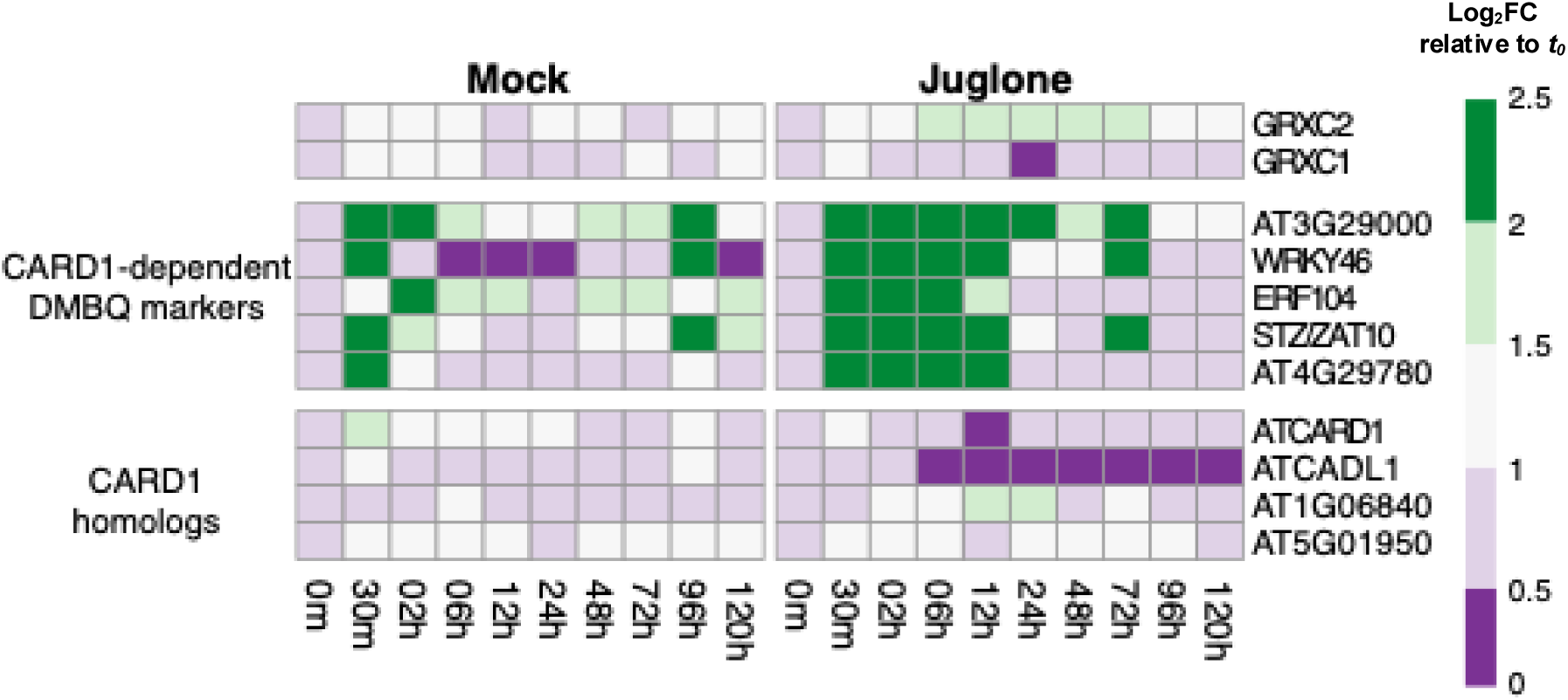
Representation of juglone-induced changes in transcript levels of genes encoding CARD1, CARD1 homologs, and CARD1-dependent 2,6-dimethoxy-1,4-benzoquinone (DMBQ) markers in Arabidopsis roots over the RNA-seq time series. Marker genes defined by Laohavisit *et al*. (2020). Color represents the log_2_fold-change (log_2_FC) in gene expression level relative to *t_0_*.

